# Endothelial Drp1 Couples VEGF-induced Redox Signaling with Glycolysis Through Cysteine Oxidation to Drive Angiogenesis

**DOI:** 10.1101/2024.06.15.599174

**Authors:** Sheela Nagarkoti, Young-Mee Kim, Archita Das, Dipankar Ash, Eric A.Vitriol, Tracy-Ann Read, Varadarajan Sudhahar, Md. Selim Hossain, Shikha Yadav, Malgorzata McMenamin, Stephanie Kelley, Rudolf Lucas, David Stepp, Eric J. Belin de Chantemele, Ruth B. Caldwell, David JR. Fulton, Tohru Fukai, Masuko Ushio-Fukai

## Abstract

Angiogenesis plays a vital role for postnatal development and tissue repair following ischemia. Reactive oxygen species (ROS) generated by NADPH oxidases (NOXes) and mitochondria act as signaling molecules that promote angiogenesis in endothelial cells (ECs) which mainly relies on aerobic glycolysis for ATP production. However, the connections linking redox signaling with glycolysis are not well understood. The GTPase Drp1 is a member of the dynamin superfamily that moves from cytosol to mitochondria through posttranslational modifications to induce mitochondrial fission. The role of Drp1 in ROS-dependent VEGF signaling and angiogenesis in ECs has not been previously described. Here, we identify an unexpected function of endothelial Drp1 as a redox sensor, transmitting VEGF-induced H_2_O_2_ signals to enhance glycolysis and angiogenesis. Loss of Drp1 expression in ECs inhibited VEGF-induced angiogenic responses. Mechanistically, VEGF rapidly induced the NOX4-dependent sulfenylation (CysOH) of Drp1 on Cys^644^, promoting disulfide bond formation with the metabolic kinase AMPK and subsequent sulfenylation of AMPK at Cys^299/304^ via the mitochondrial fission-mitoROS axis. This cysteine oxidation of AMPK, in turn, enhanced glycolysis and angiogenesis. *In vivo*, mice with EC-specific Drp1 deficiency or CRISPR/Cas9-engineered “redox-dead” (Cys to Ala) Drp1 knock-in mutations exhibited impaired retinal angiogenesis and post-ischemic neovascularization. Our findings uncover a novel role for endothelial Drp1 in linking VEGF-induced mitochondrial redox signaling to glycolysis through a cysteine oxidation-mediated Drp1-AMPK redox relay, driving both developmental and reparative angiogenesis.

## Introduction

Angiogenesis, the process of forming new blood vessels from quiescent endothelial cells (ECs), is essential for development, wound healing, and tissue repair in response to ischemic injury such as peripheral artery disease (PAD)^1^. Reactive oxygen species (ROS), particularly H_2_O_2_ derived from NADPH oxidases (NOXes) and from mitochondria at physiological levels, serve as signaling molecules that enhance vascular endothelial growth factor (VEGF)-induced signaling and angiogenic responses in ECs, as well as *in vivo* postnatal angiogenesis ^2, 3, 4, 5, 6^. However, the specific mechanisms through which diffusible H_2_O_2_ signals are transmitted to promote therapeutic angiogenesis remain elusive. ROS primarily function through the oxidation of reactive *Cys* residues, generating “*Cysteine sulfenic acid (Cys-OH)*” (*sulfenylation*), which facilitate disulfide bond formation with redox-sensitive target proteins to regulate their activity and function^7, 8^. Additionally, ECs predominantly rely on aerobic glycolysis rather than mitochondrial oxidative phosphorylation for ATP generation to drive angiogenesis^9, 10, 11, 12^. However, the mechanistic links connecting NOX-mitochondrial ROS (mitoROS)-mediated redox signaling with EC metabolism (glycolysis) in VEGF-induced angiogenesis remain to be defined.

In ECs, mitochondria serve as dynamic biosynthetic and ROS signaling organelles that contribute to EC function and vascular homeostasis^13^. Mitochondrial dynamics, governed by fusion and fission processes, adaptively maintain mitochondrial quality and function to meet the needs of the local environment^13^. Mitochondrial fusion involves Mitofusins (Mfn1 and 2) and optic atrophy 1 (OPA1) at the outer and inner membranes, respectively. Conversely, mitochondrial fission is primarily governed by Drp1 (Dynamin-related protein 1), a GTPase belonging to the dynamin family. Drp1 is recruited from the cytosol to the mitochondrial outer membrane (OMM), where it binds to Drp1 receptors such as Mff, Fis1 and MiD49 and Mid51. This interaction induces mitochondrial constriction and fragmentation in a GTP-dependent manner^13, 14, 15, 16^. Drp1 GTPase activity is regulated through various post-translational modifications (PTMs), such as phosphorylation at Ser616 (which enhances activity) or Ser637 (which inhibits activity), and S-nitrosylation at *Cys*644^14, 15, 16^. When ECs are exposed to stressors like inflammation and diabetes, Drp1 activation leads to pathological mitochondrial fragmentation (hyper fission), resulting in excessive production of mitoROS and endothelial dysfunction, including cellular senescence, and impaired angiogenesis ^13, 17, 18, 19, 20, 21, 22, 23^. However, the role of endothelial Drp1 under conditions of physiological ROS production, such as in response to VEGF signaling, and its effects on angiogenesis in ECs, as well as postnatal neovascularization *in vivo* remain unexplored.

The intricate interplay among redox signaling, mitochondria, and glycolysis during angiogenesis remains largely unexplored. To address this gap, we investigated the role of endothelial Drp1 in ROS-dependent VEGF signaling, glycolysis, and angiogenesis. This was done using various Drp1 mutant mice, including EC-specific Drp1-deficient mice and CRISPR/Cas9-engineered “redox-dead” Cys to Ala Drp1 knock-in mutant mice. Our study provides compelling evidence that VEGF rapidly induces Drp1-CysOH formation through NOX4-derived H_2_O_2_ signaling. This process drives a mitochondrial fission-mitoROS axis that promotes the cysteine oxidation of the metabolic kinase AMPK. This novel Drp1-AMPK redox relay enhances endothelial metabolism (glycolysis), which is crucial for both developmental retinal angiogenesis and reparative angiogenesis in response to tissue ischemia.

## Results

### Endothelial Drp1 is pivotal for both developmental and reparative angiogenesis in vivo

To elucidate the role of endothelial Drp1 in postnatal developmental angiogenesis, we generated conditional EC-specific Drp1 knockout (Drp1^ECKO^) mice by crossing Drp1^fl/fl^ mice with mice expressing Cre recombinase under control of the VE-cadherin promoter (Cdh5-Cre). Validation of selective Drp1 deletion in ECs was confirmed by protein analysis in primary microvascular ECs and liver tissues isolated from Drp1^ECKO^ and control (Drp1^fl/fl^ or Cdh5-Cre^+^) mice (Supplementary Fig.1a). Although global Drp1 KO mice were embryonic lethal ^24^, Drp1^ECKO^ mice were born normally and were indistinguishable from WT mice. Postnatal retina angiogenesis was evaluated in P5 pups using isolectin B4 staining. Immunofluorescence analysis demonstrated colocalization of Drp1 with isolectin B4^+^ ECs, particularly tip cells, in the P5 postnatal developmental retina (Fig.1a). Drp1^ECKO^ mice exhibited a delayed expansion of the vascular plexus toward the periphery accompanied by a significant reduction in vascular length, as well as the numbers of vascular branching points, tip cells, and filopodia in the sprouting region compared to WT mice (Fig.1b). These findings indicate that endothelial Drp1 governs vessel sprouting by facilitating tip cell filopodial extension during developmental angiogenesis.

**Figure 1.**
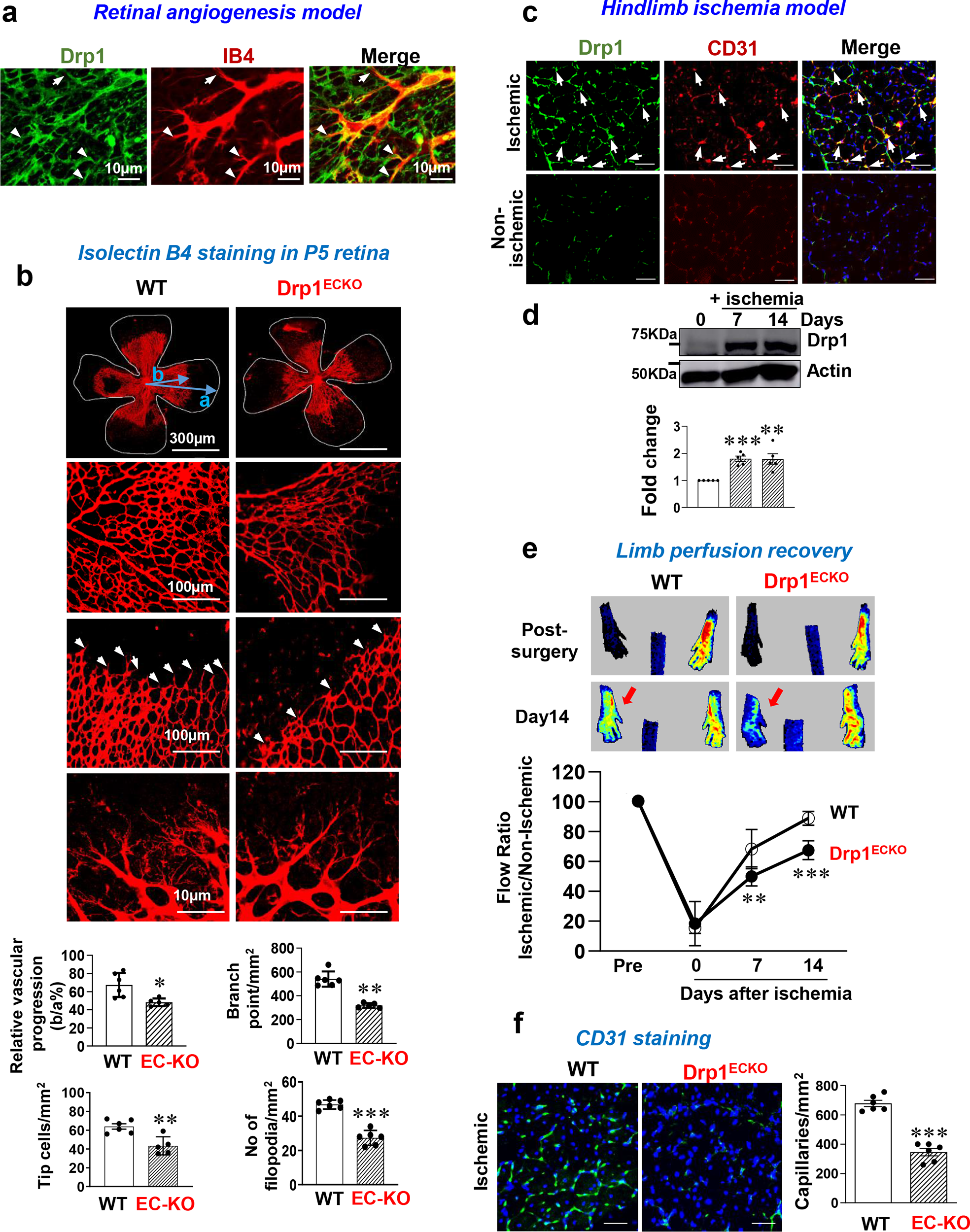
Endothelial Drp1 is essential for postnatal angiogenesis *in vivo*. **a.** Co-staining for Drp1 and isolectin B4 (IB4) on postnatal day5 (P5) mouse retina in Drp1-WT and Drp1^ECKO^ mice**. b.** Retinal whole mount staining of IB4 in P6 WT and Drp1^ECKO^ mice. The arrowheads show tip cell sprouting and filopodia. Bottom panels show quantification of vascular progression length, numbers of branching point, tip cells and filopodia. **c.** Mouse hindlimb ischemia (HLI) model: Drp1 and CD31 (EC marker) co-staining in ischemic and non-ischemic gastrocnemius (GC) muscles at day 14 post-surgery. The arrowheads show their colocalization. Scale bars=50 μm. **d.** Drp1 protein expression in ischemic GC muscles at days 0, 7 and 14 post-surgery. The bar graph shows quantification normalized by Actin. **e.** Limb perfusion recovery after HLI as determined by the ratio of foot perfusion between ischemic (left) and non-ischemic (right) legs in WT and Drp1^ECKO^ mice. Top panels show representative laser Doppler images of legs at days 0 and 14 after HLI. **F.** CD31^+^ staining (capillary density) in ischemic and non-ischemic GC muscles in WT and Drp1^ECKO^ mice on day 14. The bottom panel shows quantification. **a-f,** Data are mean ± SEM (n=5-6). *p<0.05, **p<0.005, ***p<0.001.

To explore the involvement of endothelial Drp1 in reparative angiogenesis in vivo, we employed a murine hindlimb ischemia (HLI) model, recognized as an animal model of peripheral artery disease (PAD), which induces ischemia by femoral artery ligation and excision [22,30]. Utilizing immunofluorescence (Fig. 1c) and western blotting (Fig.1d) analyses, we observed an elevation in Drp1 protein expression, which co-localized with CD31+ ECs in ischemic muscles on day 14 post-HLI. This suggests an upregulation of Drp1 in ECs during reparative angiogenesis. Furthermore, Drp1^ECKO^ mice exhibited significant reduction in limb perfusion recovery (Fig. 1e) and angiogenesis (CD31^+^ capillary density)(Fig.1f) in ischemic muscles following HLI. To confirm further the role of endothelial Drp1, we generated Tamoxifen-inducible EC-specific Drp1 KO (Drp1^iECKO^) mice by crossing Drp1^fl/fl^ mice with Cdh5-Cre^ERT^^2^ mice^25^ (Supplementary Fig.1b). Notably, ischemia-induced blood flow recovery and capillary density were significantly impaired in Drp1^iECKO^ mice (Supplementary Fig.1c and 1d). Thus, these findings suggest the crucial role of endothelial Drp1 in both developmental angiogenesis and post-ischemic reparative neovascularization *in vivo*.

### VEGF induces physiological mitochondrial fission in human ECs

We then examined underlying mechanisms using primary cultured ECs. Since Drp1 is a key regulator of mitochondrial fission, we first investigated mitochondrial shape changes in human umbilical vein ECs (HUVECs) stimulated with VEGF (20 ng/ml) which promotes EC migration, proliferation, and capillary tube formation, as previously reported^26, 27^. Figure 2a showed that in HUVECs transfected with mitochondria-targeted Mito-DsRed under overnight serum starvation, mitochondria formed a long and elongated tubular networks visualized by Zeiss confocal microscopy. VEGF stimulation predominantly induced mitochondrial fission within 30 min, which continued for at least 24 hr. Mitochondrial fission was calculated by mitochondrial fragmentation count (MFC) based on binary images showing number of mitochondrial fragments/total mitochondrial area of individual cell. To confirm this result further, we performed Tom20 staining in HUVECs visualized by Nikon spinning disk confocal microscope. The 3D binary objects were created from the deconvolved z-stacks using intensity-based thresholding (Fig. 2b) ^28^.We found that mitochondrial length was reduced in a time-dependent manner. These results suggest that VEGF stimulation at physiological concentration induces mitochondrial fission in ECs.

**Figure. 2.**
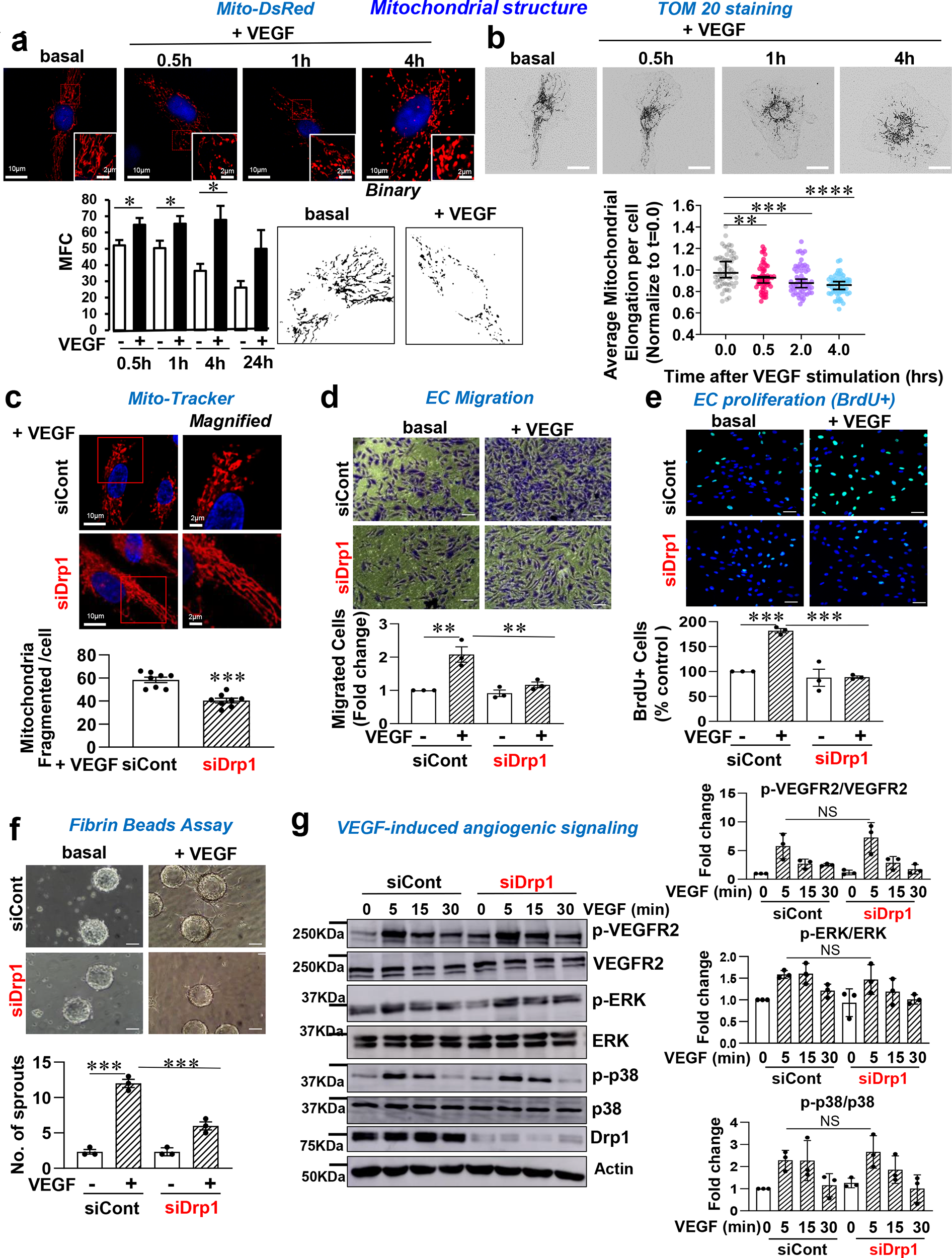
VEGF induces mitochondria fission through Drp1 that mediates VEGF-induced angiogenesis in ECs. **a.** Serum-starved HUVECs transfected with Mito-DsRed for 24 hr were stimulated with VEGF (20 ng/ml) for indicated time. Mitochondrial structure was imaged by Zeiss confocal microscopy. Bottom panels show binary images representing VEGF-induced mitochondrial fission and their quantification for mitochondrial length and mitochondrial fragmentation count (MFC). **b.** Serum-starved HUVECs were stimulated with VEGF for indicated time and mitochondrial structure was detected by TOM20 staining and visualized by Nikon CSU-W1 spinning disk confocal microscope. Bottom panel shows quantification of mitochondrial length of each cell. **c,d,e,f.** HUVECs transfected with siControl (siCont) or siDrp1 stimulated with VEGF were used to measure mitochondrial structure using Mito-Tracker staining. Bottom panel shows the number of mitochondrial fragmented cell (**c)** or EC migration (modified Boyden chamber assay)**(d)** or EC proliferation (BrdU incorporation)**(e)** or capillary tube formation in the fibrin clot using Fibrin beads assay. (scale bar=50μm)**(f)** or VEGF angiogenic signaling using Western blots with antibodies indicated **(g)**. Graph represents the averaged fold change over the basal control. Data are mean ± SEM (n=3). *p<0.05, **p<0.01, ***p<0.001. NS=not significant.

### Drp1 is involved in VEGF-induced angiogenic responses in a GTPase activity-dependent manner in ECs

We next investigated the involvement of Drp1 in VEGF-induced angiogenic responses in cultured ECs. Knockdown of Drp1 using specific siRNA in HUVEC significantly inhibited VEGF-induced mitochondrial fission (Fig. 2c), EC migration (modified Boyden chamber assay)(Fig. 2d), EC proliferation (BrdU incorporation)(Fig. 2e), and capillary tube formation (fibrin beads assay)(Fig.2f). Moreover, ECs isolated from Drp1^ECKO^ mice exhibited significant inhibition of VEGF-induced directional EC migration, assessed by the wound scratch assay compared to WT ECs (Supplementary Fig.2). Additionally, the overexpression of dominant negative Drp1-K38A (Drp1-DN) in HUVECs significantly inhibited VEGF-induced EC migration and proliferation (Supplementary Fig.3a and 3b). To address the underlying mechanism, we examined the role of Drp1 in VEGF-stimulated key angiogenic signaling in ECs. Intriguingly, Drp1 depletion with siRNA did not exert significant effects on VEGF-induced phosphorylation of VEGFR2 and its downstream signaling, p-ERK, and p-p38MAPK (Fig.2g). These findings suggest that Drp1 participates in VEGF-induced angiogenesis in a GTPase activity-dependent manner without affecting major angiogenic VEGFR2 signaling pathways.

### VEGF induces Drp1 sulfenylation at Cys^644^ through NOX4, rather than phosphorylation, which promotes Drp1 activity and angiogenesis in ECs

We next examined whether VEGF activates or upregulates Drp1 to stimulate mitochondrial fission and angiogenic responses in ECs. We found that VEGF stimulation (20 ng/ml) did not significantly alter pSer^616^–Drp1 or pSer^637^– Drp1, which regulate Drp1 activity positively or negatively, respectively, nor did it affect expression of Drp1 protein or mitochondrial dynamics-related proteins such as Mfn1 and OPA1 in ECs (Supplementary Fig.4). Since ROS, especially H_2_O_2_, derived from NOXes including NOX4, at physiological levels play a critical role in VEGF signaling and angiogenesis in ECs^29^, we next examined the role of Drp1 in VEGF-induced ROS production in ECs. We found that Drp1 siRNA had no significant effects on VEGF-induced increase in intracellular redox status, detected by CellRox (Fig. 3a). We verified that the VEGF-induced increase in CellRox fluorescence was abolished by polyethylene glycol (PEG)-conjugated catalase or NOX4 siRNA (data not shown), indicating that its signal mainly reflects Nox4-derived intracellular H_2_O_2_ levels.

**Figure 3.**
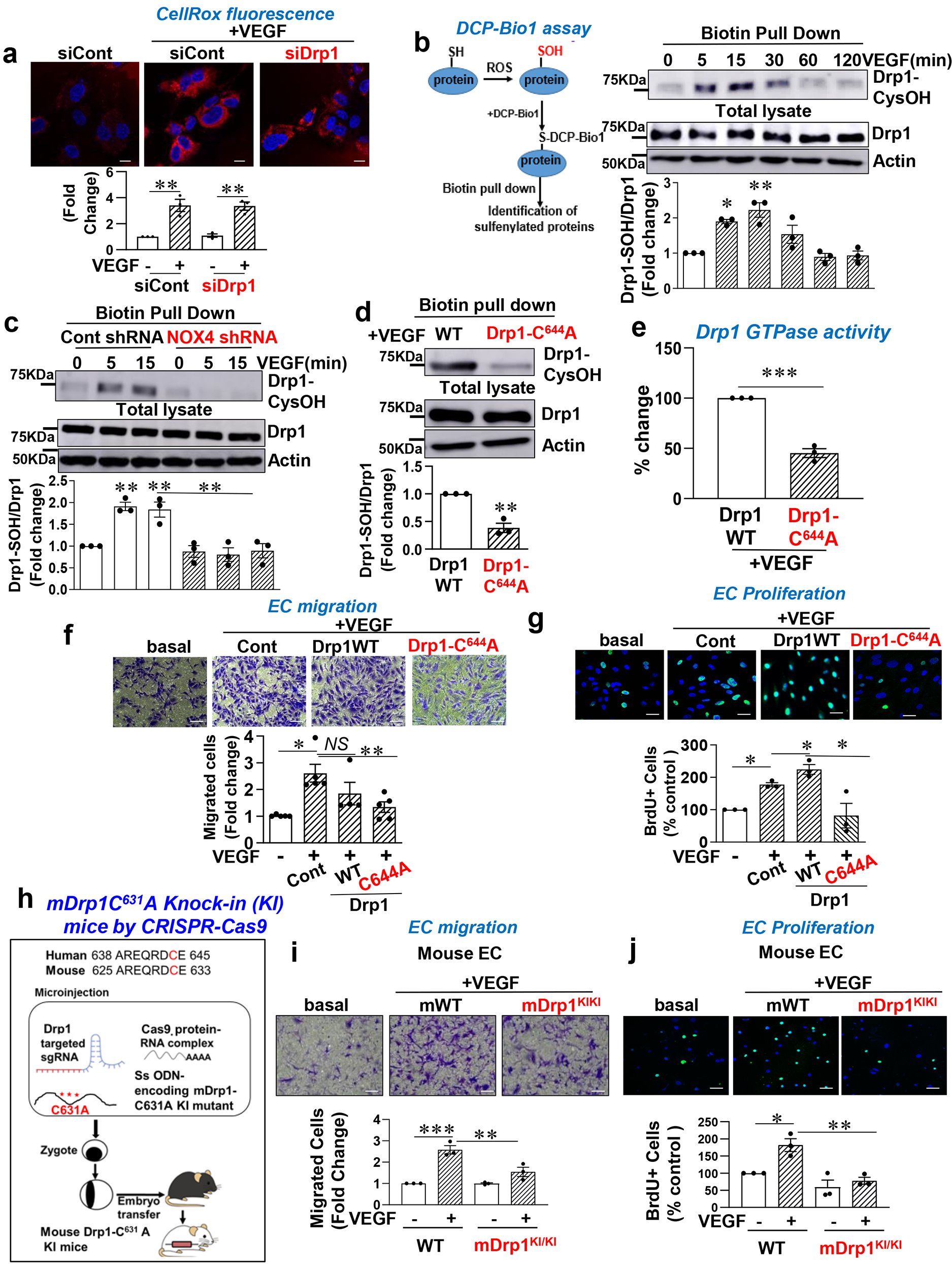
VEGF induces sulfenylation of Drp1 at Cys^644^ in a Nox4-dependent manner, which promotes mitochondrial fission, mitoROS, and angiogenesis in ECs. **a.** HUVEC transfected with control or Drp1 siRNAs stimulated with VEGF (20ng/ml) for 5 min were used to measure CellRox fluorescence and DAPI staining (blue). The bottom panel represents the averaged fold change of fluorescence intensity over the basal control. **b,c,d.** DCP-Bio1 assay: Left panel, Scheme of DCP-Bio1 assay to detect sulfenylated protein. Right panel, HUVECs were stimulated with VEGF for indicated time **(b)**. HUVECs transduced with Ad.null (Cont) shRNA or Ad.shNox4 were stimulated with VEGF for indicated time **(c)**. HUVECs transduced with Ad.Drp1-WT or Ad.Drp1-C^644^A were stimulated with VEGF for 5 min **(d)**. DCP-labeled lysates were pulled down with streptavidin beads and then immunoblotting (IB) with Drp1 antibody (Ab). Bottom panels represent the averaged fold-change of Drp1-CysOH/Drp1. **e.** HUVECs transduced with Ad Drp1-WT or Ad.Drp1-C^644^A were used to measure Drp1GTPase activity. **f and g.** HUVECs transduced with Ad. Cont or Ad.Drp1-WT or Ad.Drp1-C^644^A were used to measure VEGF-induced EC migration (modified Boyden chamber assay)**(f)** or EC proliferation (BrdU incorporation)**(g)**. **h.** Schematic diagram of CRISPR-generated “redox dead” mDrp1-C^631^A (corresponding to human Drp1-C^644^A) knock-in (KI)(mDrp1^KI/KI^ mice. CRISPR sgRNA targeting the mDrp1 gene, Cas9 mRNA, and single stranded oligo donor DNA (ssODN) encoding Drp1 mutations was injected into mouse zygotes. The precise mutation of mDrp1^KI/KI^ mice was verified by combination of PCR and a next generation sequencing. **i and j.** Aortic ECs isolated from WT or mDrp1^KI/KI^ mice were used to measure VEGF-induced EC migration (modified Boyden chamber assay)**(i)** or EC proliferation (BrdU incorporation)**(j)**. **a-j,** Data are mean ± SEM (n=3). *p<0.05, **p<0.01, ***p<0.001.

We next investigated whether Drp1 is a downstream target of VEGF-induced H_2_O_2_ to promote EC angiogenesis. Since ROS induce the oxidation of Cys residues in target proteins to form CysOH (sulfenylation), and Drp1 has a crucial redox-sensitive Cys^644^ residue^30^, we examined if VEGF induces CysOH formation of Drp1 in ECs. Using a biotin-conjugated CysOH trapping probe, DCP-Bio1, we observed that VEGF stimulation rapidly increased Drp1-CysOH formation within 5 min, peaking at 15 min, and gradually returning to basal levels within 60 min (Fig.3b). Notably, this VEGF-induced Drp1 sulfenylation was abolished by Nox4 shRNA (Fig.3c), which inhibited VEGF-induced H_2_O_2_ production and angiogenic responses in ECs^29^. To assess the role of Cys^644^ in VEGF-induced Drp1-CysOH formation, we engineered adenovirus expressing Drp1-C^644^A, where Cys^644^ is substituted with Ala^644^. VEGF-induced CysOH formation of Drp1 was significantly inhibited by Drp1-C^644^A compared to Drp1-WT (Fig. 3d). Importantly, the overexpression of Drp1-C^644^A significantly inhibited VEGF-induced Drp1GTPase activity (Fig. 3e). These findings imply that VEGF induces sulfenylation of Drp1 at Cys^644^ in a NOX4-derived H_2_O_2_, thereby activating Drp1 in ECs.

We next investigated the functional role of Drp1 sulfenylation or Drp1 activation in VEGF-induced angiogenesis in ECs. Overexpression of Drp1-C^644^A, or Drp1-DN significantly inhibited VEGF-induced EC migration (Fig.3f, Supplementary Fig.3a) and EC proliferation (Fig.3g, Supplementary Fig.3b) compared to control or Drp1-WT. To further underscore the functional significance of endogenous Cys^644^ oxidation of Drp1, we generated “redox-dead” hCys644 (mouse mCys631) to Ala Drp1 knock-in (KI) mutant (mDrp1-KI) mice using CRISPR-Cas9 genome editing (Fig.3h). ECs isolated from mDrp1-KI mice (mDrp1-KI ECs) exhibited marked inhibition of VEGF-induced EC migration (Fig.3i) and EC proliferation (Fig.3j) compared to WT-ECs. Taken together, these findings suggest that VEGF-induced sulfenylation of Drp1 at Cys^644^ is required for VEGF-induced angiogenic responses.

### VEGF-induced sulfenylation of Drp1 at Cys^644^ facilitates mitochondrial fission and mitoROS production, which promotes glycolysis

Given that Drp1-mediated mitochondrial fission is known to increase mitoROS production^17^ and mitoROS is essential for VEGF-induced angiogenesis ^29^, we investigated the impact of Cys^644^ oxidation of Drp1 on VEGF-induced mitochondrial fission and mitoROS production. Overexpression of Drp1-C^644^A, but not Drp1-WT, significantly inhibited VEGF-induced mitochondrial fission, as assessed by Mito-DsRed staining (Fig. 4a), and mitoROS production, as measured by Mito-SOX fluorescence (Fig. 4b). Given that glycolysis serves as the primary source of ATP driving angiogenesis in ECs, we examined whether Drp1 Cys oxidation is involved in VEGF-induced glycolysis in ECs by analyzing the extracellular acidification rate (ECAR), a measure of glycolytic flux, using the XF24 extracellular flux analyzer. Figures 4c and 4d showed that overexpression of Drp1-C^644^A or mitochondria-targeted catalase (Mito-catalase) significantly inhibited VEGF-induced glycolysis. Of note, Mito-catalase overexpression had no effects on VEGF-induced Drp1-CysOH formation (data not shown). These findings suggest that VEGF-induced sulfenylation of Drp1 mediated through NOX4-derived H_2_O_2_, stimulates mitochondrial fission and mitoROS (H_2_O_2_) production, thereby promoting glycolysis and angiogenesis in ECs (Fig. 4e).

**Figure 4.**
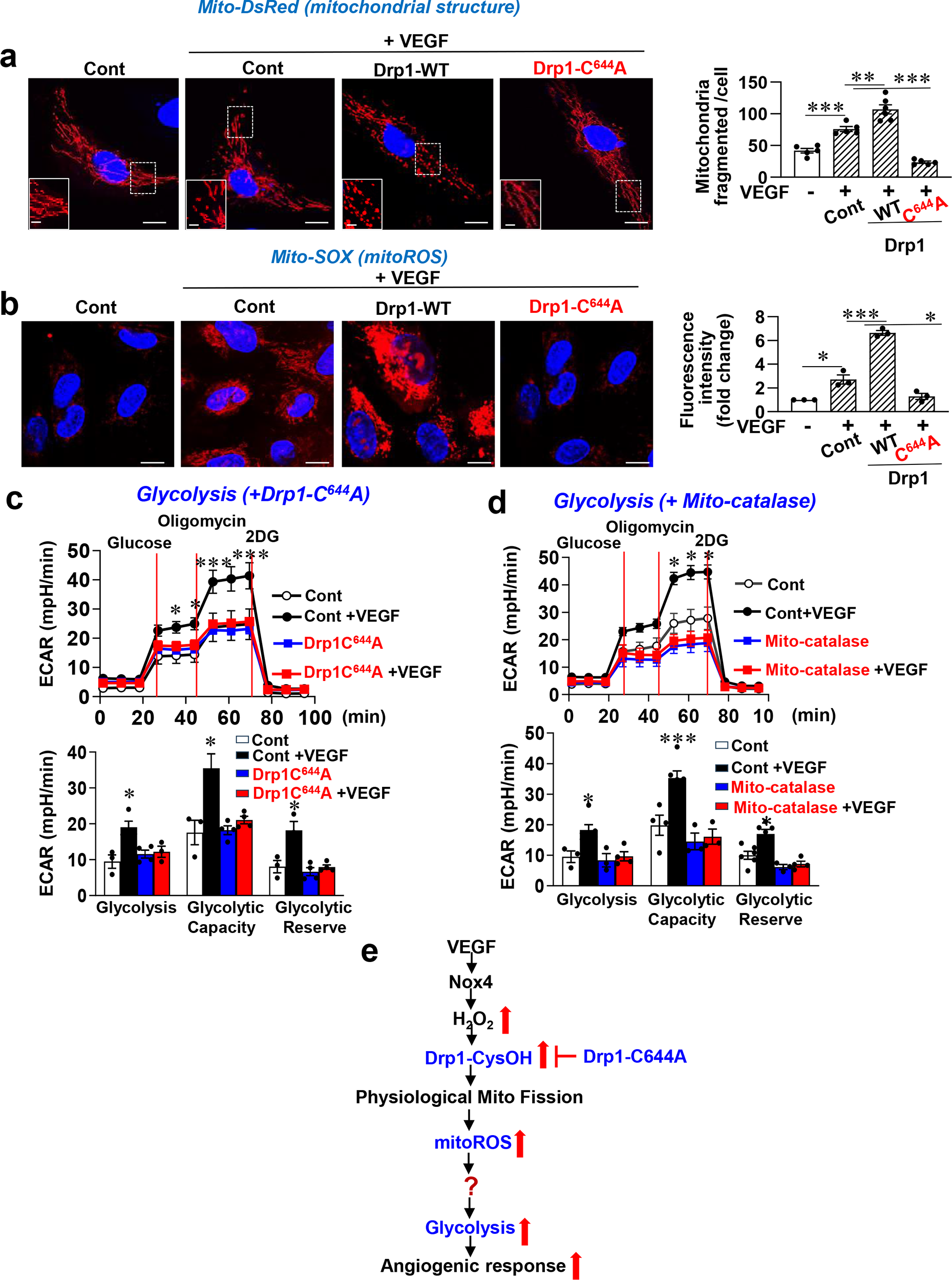
VEGF-induced sulfenylation of Drp1 promotes mitochondrial fission, mitoROS and glycolysis in ECs. **a and b.** HUVECs transduced with Ad.Cont, Ad.Drp1-WT, Ad.Drp1-C^644^A were transfected with Mito-DsRed **(a)** or incubated with Mito-SOX **(b)**, and then stimulated with VEGF for 30 min to measure mitochondrial fission and mitoROS production, respectively. (scale bar=10mm) **c and d.** HUVECs transduced with Ad.Cont, Ad.Drp1-C^644^A **(c)** or Ad.Mito-catalase **(d)** were stimulated with VEGF for 1 hr, followed by measuring glycolysis (ECAR) using Seahorse analyzer. Data are mean ± SEM (n=3). *p<0.05, **p<0.01, ***p<0.001. **e**. Schematic model: VEGF induces Drp1-CysOH formation at Cys644 via Nox4-derived H2O2, which promotes physiological mitochondrial fission-mitoROS axis that enhances glycolysis and angiogenesis. Question mark (?) represents targets of mitoROS involved in glycolysis and angiogenesis.

### VEGF-induced sulfenylation of Drp1 at Cys^644^ is required for disulfide bond formation with AMPK and activation of AMPK in ECs

To elucidate the mechanism by which Drp1-CysOH promotes glycolysis, we performed preliminary studies to identify the Drp1 binding partners known to regulate glycolysis. Co-immunoprecipitation (Co-IP) analysis revealed that Drp1 binds to AMPK, a pivotal angiogenic and metabolic kinase, upon VEGF stimulation (Fig. 5a). Moreover, non-reducing gel analysis demonstrated that VEGF stimulation induces disulfide bond formation between Drp1 and AMPK (Fig. 5b). To validate the role of Drp1-Cys^644^OH in redox-sensitive Drp1-AMPK complex formation, we performed Bimolecular Fluorescence Complementation (BiFC) assay. Figure 5c shows that co-transfection of AMPK fused to the Venus N terminus (VN-AMPK) and Drp1 fused to the Venus C terminus (VC-Drp1-WT) induced yellow fluorescent protein (YFP) fluorescence in H_2_O_2_-stimulated cells. Conversely, cells transfected with VN-AMPK with a scrambled peptide fused to VC (VC-peptide, negative control) or Drp1-C^644^A mutant fused to the VC (VC-Drp1-C^644^A) showed no fluorescence (Fig. 5c). We confirmed that VN-AMPK Co-IP with VC-Drp1-WT, but not with VC-Drp1-C^644^A (Fig. 5d). We then examined whether Drp1 sulfenylation is necessary for VEGF-induced AMPK activity which is regulated by various post-translational modifications such as phosphorylation. Figure 5e demonstrated that overexpression of Drp1-C^644^A had no significant effects on VEGF-induced AMPK phosphorylation (p-AMPK) but markedly inhibited p-ACC, a direct downstream substrate of AMPK reflecting its activation. These findings suggest that VEGF induced formation of Drp1-Cys^644^OH is indispensable for Drp1-AMPK disulfide bond formation and subsequent AMPK activation (Fig. 5f).

**Figure 5.**
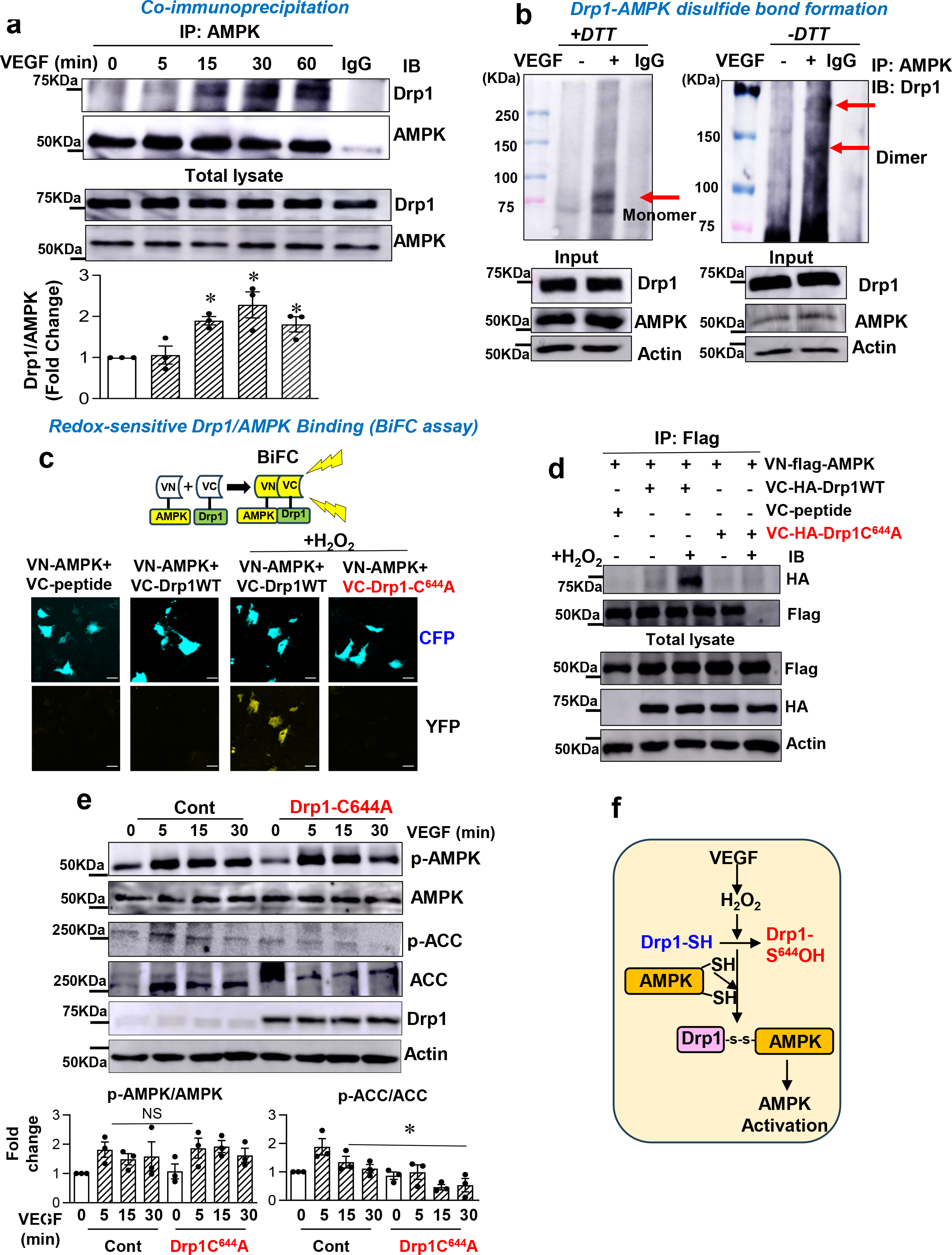
VEGF-induced sulfenylation of Drp1 at Cys^644^ is required for disulfide bond formation with AMPK and activation of AMPK in ECs. **a**. HUVECs stimulated with VEGF (20ng/ml) for indicated times were IP with AMPK Ab or IgG (negative control), followed by IB with anti-Drp1 or AMPK Ab. **b**. HUVECs stimulated with VEGF for 15 min were IP with AMPK Ab or IgG, followed by SDS-PAGE under nonreducing (-DTT) and reducing (+DTT) conditions and IB with anti-Drp1Ab. **c and d**. BiFC assay: Cos1 cells were co-transfected with N-terminal Venus (VN)-flag-AMPK (VN-flag-AMPK) and C-terminal Venus (VC)-negative peptide (VC-peptide) or VC-HA-Drp1-WT or VC-HA-Drp1-C^644^A. Cells of each group stimulated with H2O2 (200 mM) for 10 min were imaged by confocal microscopy. (scale bar=20mm) **(c)** or IP with anti-Flag Ab, followed by IB with anti-HA or anti-Flag Ab **(d)**. In **(c)**, The YFP (Yellow) fluorescence shows interaction of AMPK1 and Drp1. **e**. HUVECs transduced with Ad.Cont or Ad.Drp1-C^644^A were stimulated with VEGF for indicated times, and then used for IB with Abs indicated. **f.** Schematic model: VEGF-induced Drp1-Cys^644^OH promotes disulfide formation with AMPK-SH, followed by AMPK activation. Data are mean ± SEM (n=3). *p<0.05, **p<0.01, ***p<0.001.

### VEGF-induced sulfenylation of Drp1 at Cys^644^ is required for mitoROS-dependent AMPK Cys oxidation, driving glycolysis and angiogenesis in ECs

Given that H_2_O_2_ stimulation is shown to enhance AMPK activity via S-glutathionylation at Cys299/304 ^31^, we examined whether VEGF stimulation induces Cys oxidation of AMPK in ECs. VEGF stimulation led to a significant increase in CysOH formation of AMPK within 15 min (Fig. 6a), which was inhibited by overexpression of Drp1-C^644^A (Fig. 6b), Mito-catalase (Fig. 6c) or AMPK-C^299/304^A (Fig. 6d). These results indicate that VEGF-induced Drp1-Cys^644^OH-mediated mitoROS production promotes AMPK oxidation at Cys^299/304^. Notably, this was associated with VEGF-stimulated translocation of Drp1 and AMPK to mitochondrial fraction (Supplementary Fig. 5). We next assessed the role of AMPK sulfenylation in VEGF-induced glycolysis and angiogenic responses. Overexpression of AMPK-C^299/304^A significantly inhibited VEGF-induced glycolysis (ECAR)(Fig.6e), as measured by Seahorse assay, as well as EC migration (Fig.6f) and EC proliferation (Fig.6g). Thus, these results suggest that VEGF-induced Drp1-Cys^644^OH promotes its translocation to the mitochondria by forming disulfide bond with AMPK, which facilitates the oxidation and activation of AMPK by mitoROS, thereby driving glycolysis and angiogenesis in ECs (Fig. 6h).

**Figure 6.**
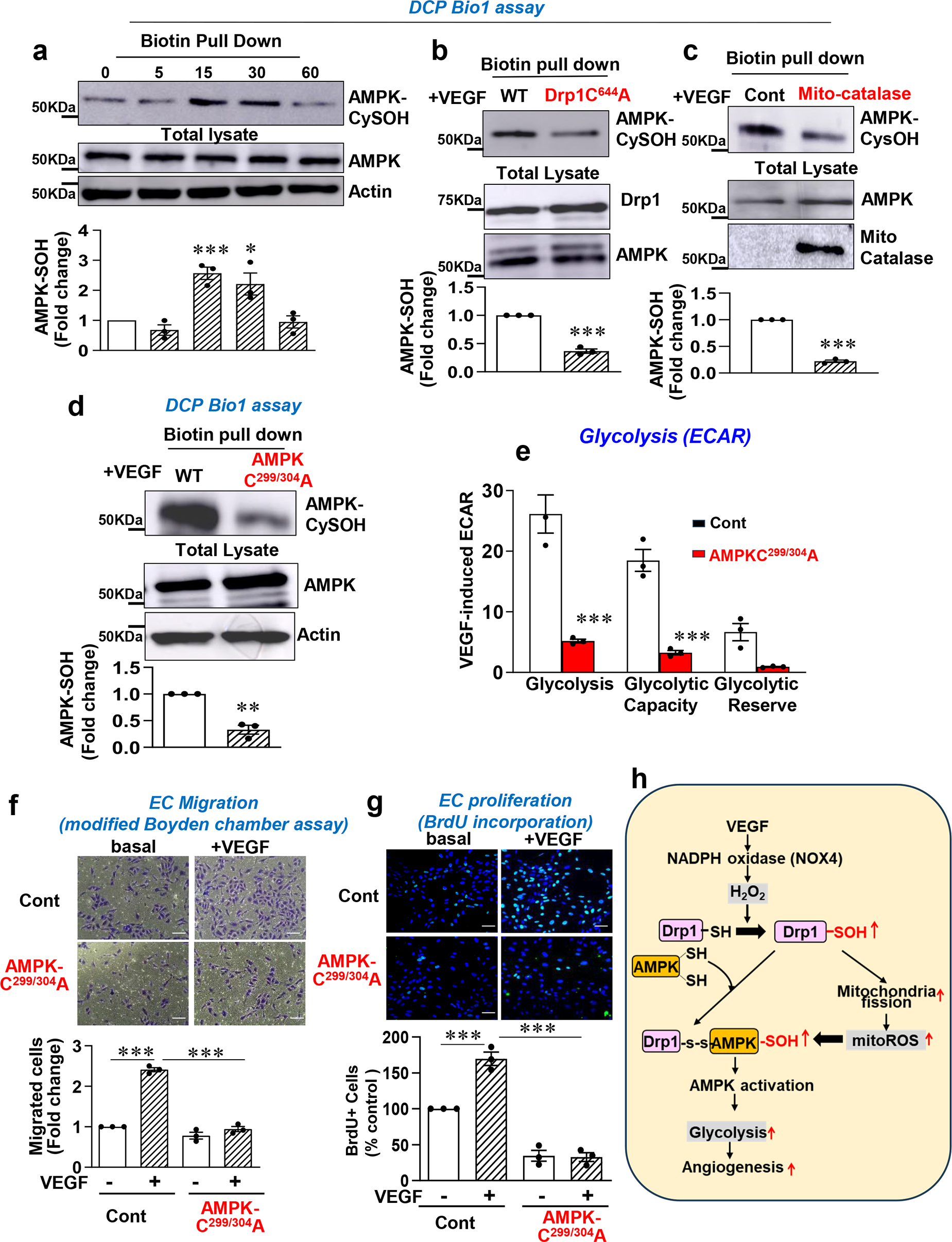
VEGF-induced sulfenylation of Drp1 at Cys^644^ and mitoROS are required for Cys oxidation of AMPK, which promotes angiogenic responses in ECs. **a,b,c,d.** HUVECs were stimulated with VEGF (20ng/ml) for indicated time **(a)**. HUVECs transduced with Ad.Drp1-WT or Ad.Drp1-C^644^A (**b**) or transduced with Ad.Cont or Ad.Mito-catalase (**c**) or transduced with Ad.AMPK-WT or Ad.AMPK-C^299/304^A (**d**), were stimulated with VEGF for 15min. DCP-Bio1-labeled lysates were pulled down with streptavidin beads, followed by IB with AMPK Ab. Graphs represent the averaged fold change of AMPK-CysOH/AMPK. **e.** HUVECs transduced with Ad.Cont or Ad.AMPK-C^299/304^A stimulated with VEGF for 1 hr were used to measure glycolysis (ECAR) using Seahorse analyzer. **f and g**. HUVECs transduced with Ad.Cont or Ad.AMPK-C^299/304^A were used to measure VEGF-induced EC migration (modified Boyden chamber assay)**(f)** or EC proliferation (BrdU assay)**(g)**. Lower panel represents the averaged fold change from the basal control. Data are mean ± SEM (n=3). *p<0.05, **p<0.01, ***p<0.001. **h.** Schematic model: VEGF stimulation induces NOX4-dependent sulfenylation of Drp1 at Cys^644^, which facilitates the formation of disulfide with AMPK-SH, leading to translocation from cytosol to mitochondria, subsequently activating AMPK via oxidation at Cys^299/304^ by mitochondria fission-mitoROS axis.

### Drp1-C^644^A (mDrp1-C^631^A) knock-in (mDrp1-KI) mice exhibit impaired developmental and reparative angiogenesis *in vivo*

To determine the role of endogenous Drp1 Cys oxidation in developmental and reparative angiogenesis *in vivo*, we used the Cys^644^ oxidation-defective “redox-dead” mDrp1-C^631^A-KI mutant (mDrp1-KI) mice, as detailed in Figure 3h. Homozygous mDrp1^KI/KI^ mice displayed no phenotype during embryogenesis, exhibiting viability and normal body weight compared to WT littermate. In a postnatal developmental retinal angiogenesis model, mDrp1^KI/KI^ mice exhibited delayed expansion of the vascular plexus toward the periphery, along with a significant reduction in vascular length and the numbers of vascular branching points, tip cells, and filopodia in the sprouting region compared to WT mice (Fig. 7a). Using the hindlimb ischemia PAD model, we observed a marked reduction of Drp1 CysOH formation (Fig. 7b) and blood flow recovery (Supplementary Fig. 6) in ischemia muscle of mDrp1^KI/KI^ mice. Furthermore, to eliminate the contribution of bone marrow (BM) cells, we performed BM transplantation (BMT) experiments. Lethally irradiated mDrp11^KI/KI^ mice transplanted with WT-BM versus WT mice reconstituted with WT-BM exhibited significant impairment of limb perfusion recovery (Fig.7c) and CD31+ capillary density in ischemic muscle (Fig. 7d). These findings suggest that Drp1 sulfenylation at Cys^644^ in tissue-resident cells including ECs is crucial for ischemia-induced reparative neovascularization.

**Figure 7.**
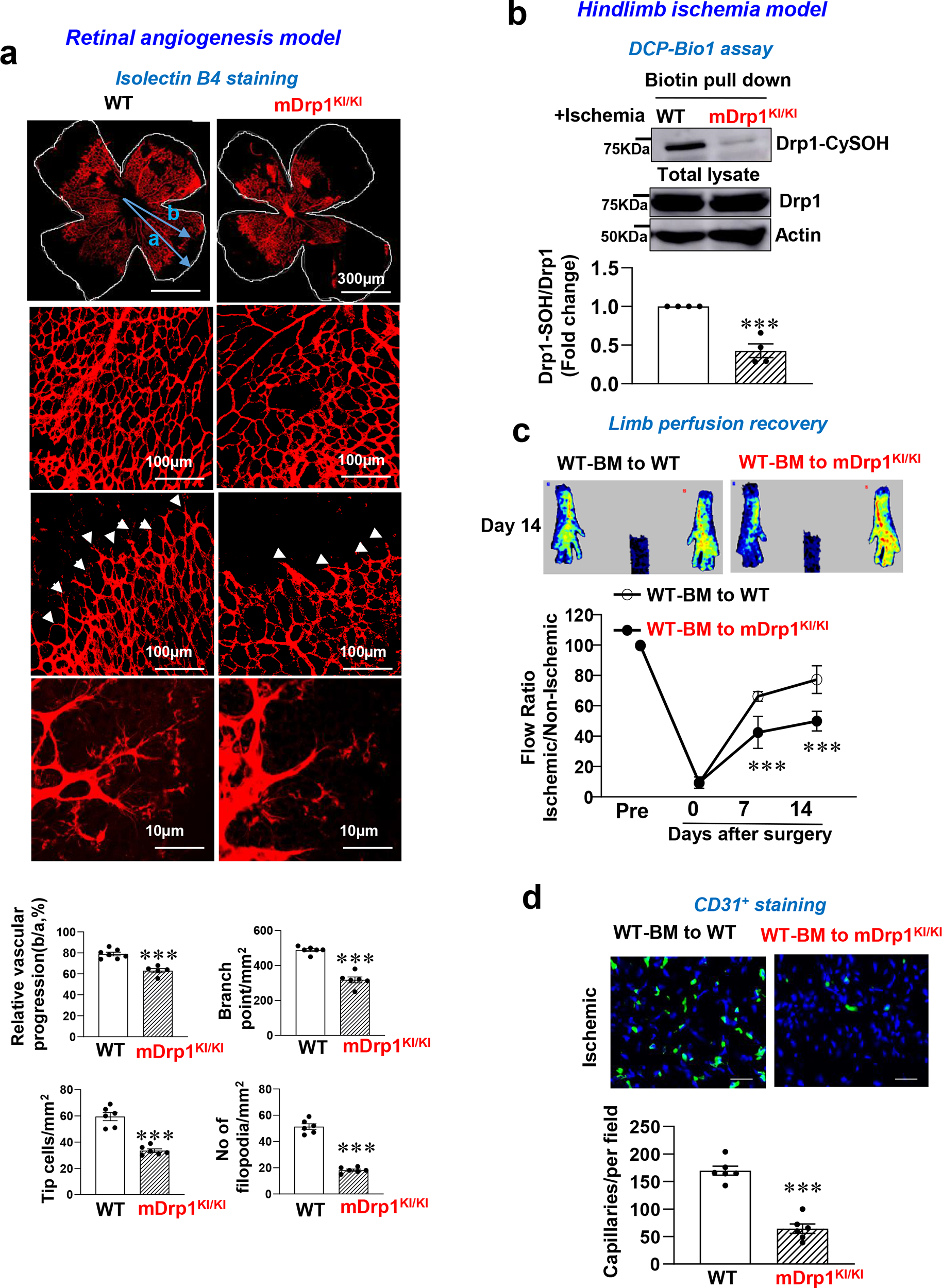
Drp1 sulfenylation at Cys^644^ is required for developmental angiogenesis and reparative neovascularization *in vivo*. **a.** Retinal angiogenesis model: Retinal whole mount staining of isolectin B4 in P5 WT and CRISPR-generated “redox dead” mDrp1^KI/KI^ mice. The arrowheads show tip cell sprouting and filopodia. **b,c,d.** HLI model: WT or mDrp1^KI/KI^ mice were subjected to HLI, and tibialis anterior (TA) ischemic muscle was isolated to measure Drp1-CysOH formation using DCP-Bio1 (**b**). **c and d**. Irradiated WT or mDrp1^KI/KI^ mice were transplanted with bone marrow (BM) from WT mice to eliminate the contribution of mDrp1^KI/KI^ myeloid cells. After 6 weeks of BM transplantation (BMT), limb perfusion recovery after HLI was determined by the ratio of foot perfusion between ischemic and non-ischemic legs using a laser Doppler flow analyzer. Top panels show representative laser Doppler images of legs on day 14 after HLI **(c)**. CD31^+^ capillary density (angiogenesis) in ischemic GC muscles on day 21 after HLI. Scale bars=5 μm. The bottom panel shows quantification (**d**). Data are mean ± SEM (n=6) ***p<0.001.

## Discussion

The interplay among redox signaling, mitochondria, and glycolysis during angiogenesis remains poorly understood. Our study reveals a previously unrecognized role of endothelial Drp1 as a redox sensor that transmits VEGF-induced H_2_O_2_ signal to enhance aerobic glycolysis through oxidation at Cys^644^, thereby promoting developmental and reparative angiogenesis. Mechanistically, VEGF stimulation triggers mitochondrial fission by increasing NOX4-dependent sulfenylation of Drp1 at Cys^644^. This process facilitates the formation of disulfide bonds between Drp1 and AMPK, promoting translocation to mitochondria and the increased proximity to mitoROS activates AMPK via Cys oxidation. Thus, this unexpected redox signaling cascade links Drp1 with AMPK (redox relay) to orchestrate aerobic glycolysis and angiogenic responses in ECs, contributing to postnatal developmental and reparative angiogenesis *in vivo*.

ECs predominantly rely on glycolysis, rather than mitochondrial oxidative phosphorylation, as a major source of ATP to drive angiogenesis^9, 32^, ^33^. Under physiological conditions, mitochondrial dynamics are tightly regulated by the balance of fission and fusion, which maintains normal mitochondrial function ^13^. However, the specific role of the endothelial mitochondrial fission protein Drp1 in postnatal developmental and reparative angiogenesis *in vivo* has not been reported. It has been shown that global Drp1 deficient mice exhibit early embryonic lethality (embryonic days 11.5) ^24^. In the present study, EC-specific Drp1 KO mice reveal that loss of Drp1 in ECs does not result in an overt phenotype such as embryonic lethality but instead we observed impaired vessel sprouting during developmental angiogenesis in the retina as well as compromised reparative neovascularization following ischemic injury. To the best of our knowledge, this is the first evidence showing an essential role of endothelial Drp1 in angiogenesis utilizing genetically modified mice deficient in Drp1 only in ECs. In contrast to our finding, it has been reported that EC-specific knockout of the inner mitochondrial fusion protein, OPA1, in mice resulted in impaired pathological tumor angiogenesis due to enhanced NFkB activation, independent of its better known function mediating mitochondrial fusion^34^. Although the role of endothelial Drp1 in pathological angiogenesis (such as tumor angiogenesis and diabetic retinopathy) needs to be explored, these findings support the notion that both the mitochondrial fission protein, Drp1 and the mitochondrial fusion protein, OPA1 play important roles regulating postnatal angiogenesis *in vivo* albeit through distinct mechanisms.

A specific role for endothelial Drp1 in mediating VEGF signaling and angiogenic responses in ECs has not been reported. In current study, we used confocal microscopy and Nikon spinning disk confocal microscopy to image live cells at subcellular resolution ^28^. This revealed that serum starved HUVEC displayed an elongated filamentous mitochondrial network in their basal state. Upon VEGF stimulation, rapid mitochondrial fission became pronounced within 30 min and persisted for at least 4 hrs. Drp1 depletion, overexpression of a dominant negative (DN)-Drp1 (which inhibits Drp1GTPase activity), or ECs isolated from Drp1^ECKO^ mice, exhibited impaired VEGF-induced EC migration, proliferation, and capillary tube formation, underscoring a critical role of Drp1 in VEGF-induced angiogenesis. Conversely, under pathological conditions such as inflammation and diabetes, where the balance between fusion and fission is disrupted, hyperactivation of Drp1 and excessive mitochondrial fission occur, leading to oxidative stress, EC dysfunction including senescence^18,35^ and angiotensin II-induced pathologies ^36^. These data highlight the ability of Drp1 to have either beneficial or harmful actions depending on the extent of Drp1 activation, the presence of stimuli such as growth factors or inflammatory cytokines, and changes in the microenvironment such as hypoxia or hyperglycemia. However, we cannot eliminate the possibility that previous studies showing deleterious effects of mitochondrial division inhibitor 1 (Mdivi1), a small molecule inhibitor of Drp1, might be due to non-specific effects from inhibition of mitochondrial complex I ^37^. Additionally, it has been reported that higher concentrations of VEGF (50 ng/ml) can induce mitochondrial fusion and increased expression of the inner mitochondrial fusion protein, OPA1, in ECs and that OPA1 depletion inhibits VEGF-induced angiogenesis by activating NFkB in a manner independent of mitochondrial fusion^34^. Similarly, knockdown of the outer mitochondrial fusion protein, MFNs, blocked VEGF-induced angiogenesis by reducing expression of respiratory chain components ^35^. Accordingly, the balance between mitochondrial fission and fusion can be influenced by the VEGF concentration, the cell types, and culture conditions. Thus, it is likely that imbalance between mitochondrial fission and fusion impairs VEGF-induced angiogenesis and that both processes may play crucial roles in this response.

The activity of Drp1 GTPase is regulated by various PTMs such as phosphorylation or S-nitrosylation at Cys^644^ or other modification^14, 15, 16^. In our study, VEGF did not phosphorylate Drp1 at Ser^616^ or Ser^637^ or increase Drp1 expression in ECs. Instead, experiments employing DCP-Bio1 assay, Drp1-C^644^A mutant, which inhibits Drp1-GTPase activity, or ECs isolated from redox-dead” mDrp1-C^631^A^KI/KI^ mutant mice revealed that VEGF rapidly induced the sulfenylation of Drp1 at Cys^644^ in a NOX4-dependent manner, which promoted physiological mitochondrial fission, mitoROS production, and angiogenic responses in ECs. Notably, in quiescent ECs that have a reduced redox state, Drp1 exists in a reduced form (Drp1-SH) and inactive state, bound to the thiol reductase protein disulfide isomerase A1 (PDIA1) ^38^. Therefore, reduced Drp1-SH in ECs at baseline is required for VEGF-induced Drp1-CysOH formation by Nox4-derived H_2_O_2_, which promotes a physiological mitochondrial fission-mitoROS axis that enhances angiogenic responses. Furthermore, the physiological induction of Drp1 sulfenylation by VEGF can be distinguished from the pathological excessive sulfenylation/activation of Drp1 observed in basal quiescent ECs lacking thiol reductase PDIA1 or under diabetic conditions that results in hyper mitochondrial fission and EC senescence ^38^. Further studies are required to identify other redox-sensitive Cys residues on Drp1 that are substrates for VEGF-induced sulfenylation and to determine whether other-post-translational modifications of Drp1, such as S-glutathionylation or S-nitrosylation or S-sulhydration, play a role in VEGF-induced angiogenesis in ECs. The *in vivo* significance of Drp1-Cys^644^ oxidation was demonstrated by experiments in mice harboring a ‘redox-dead” mDrp1-C^631^A^KI/KI^ mutant that revealed impaired developmental retinal angiogenesis and reparative angiogenesis in response to ischemia. Additionally, BM transplant experiments where irradiated mDrp1-C^631^A^KI/KI^ mice were reconstituted with bone marrow from Drp1 WT BM support the concept that Drp1 Cys^644^ oxidation in tissue resident cells including ECs play a critical role in postnatal reparative angiogenesis.

Angiogenesis in ECs is a highly energy-demanding process that primarily relies on glycolysis, instead of mitochondrial oxidative phosphorylation^39^. However, the role of mitochondrial dynamic proteins in regulating glycolysis in ECs remains poorly understood. Our study establishes a mechanistic connection between NOX-mitoROS-mediated redox signaling and glycolysis through Drp1 sulfenylation in VEGF-induced angiogenesis. Unexpectedly, we found that overexpression of the Drp1-C^644^A mutant or Mito-catalase significantly inhibited VEGF-induced glycolysis in ECs. This suggests that Drp1 sulfenylation at Cys^644^ via NOX4-derived H_2_O_2_, along with the enhanced activity of the mitochondrial fission-mitoROS axis, is essential for VEGF-induced glycolysis in ECs. Mechanistically, we identified AMPK, a key regulator of metabolism and angiogenesis, as a novel redox-sensitive binding partner of Drp1. Using Co-IP, BiFC, and mitochondrial fractionation, our experiments indicate that VEGF-induced sulfenylation of Drp1 at Cys^644^ promotes its binding to AMPK-SH, thereby forming disulfide bond between Drp1-AMPK. This interaction facilitated mitochondrial localization of AMPK, resulting in cysteine oxidation (sulfenylation) and activation by mitoROS (Fig. 6h). Previous studies have shown that exogenous H_2_O_2_ increases AMPK activity through oxidative modification of Cys299/304 ^31^. Consistently, we found that overexpression of AMPK-C^299/304^A resulted in the inhibition of VEGF-induced AMPK sulfenylation, glycolysis and angiogenic responses in ECs. These findings suggest that Drp1 sulfenylation at Cys^644^ by Nox4-derived H_2_O_2_ promotes the mitochondrial localization and sulfenylation of AMPK at Cys^299/304^, thereby enhancing glycolysis and angiogenesis. Identifying the Cys residue of AMPK forming a disulfide bond with Drp1-Cys^644^OH, as well as other downstream targets of sulfenylated Drp1 that might contribute to enhanced glycolysis and angiogenesis will be the subjects of future studies.

Collectively, our findings advance the understanding of how VEGF-induced redox signals are sensed and converted into signals that enhance glycolysis and angiogenesis. Additionally, uncover an unexpected role for the mitochondrial fission protein Drp1 as a redox sensor that facilitates the transmission of NOX4-derived H_2_O_2_ signals via sulfenylation of Cys^644^. This mechanism activates the mitochondrial fission-mitoROS axis, leading to the oxidative activation of the key metabolic enzyme AMPK through the formation of a disulfide bond between Drp1 and AMPK in ECs. Consequently, this process enhances endothelial glycolysis, promoting developmental and reparative angiogenesis *in vivo*. Furthermore, our study highlights a novel cysteine oxidation-mediated Drp1-AMPK redox relay that coordinates VEGF-induced redox signals and enhanced aerobic glycolysis, offering a potential therapeutic target for promoting angiogenesis. This discovery may be of significant therapeutic utility in developing new treatments for ischemic cardiovascular diseases.

## Materials and Methods

### Animal Study

The use of mice was in accordance with the National Institutes of Health Guide for the Care and Use of Laboratory Animals and relevant ethical regulations. The animal protocol used in this study was approved by the institutional Animal Care committee and institutional Biosafety Committee of Augusta University.

### Generation of conditional EC-specific Drp1 KO (Dp1^ECKO^) mice and inducible EC-specific Drp1 KO (Drp1^iECKO^) mice

Dp1^ECKO^ mice were generated by crossing Drp1^fl/fl^ mice ^24^ with VE-Cadherin (Cdh5)-Cre mice on a C57BL/6J background and used at 8-12 weeks. Tamoxifen-inducible Drp1^iECKO^ mice were generated by crossing Drp1^fl/fl^ mice with Cdh5-Cre^ERT^^2^ mice^40^ on a C57BL/6J background. To induce postnatal deletion of endothelial Drp1, we administered tamoxifen (50 mg kg−1 of body weight) by intraperitoneal (i.p.) injection into control, wild type (WT)(Drp1^fl/fl^ or Cdh5-Cre negative) mice, or Drp1^fl/fl^/Cdh5-Cre^ERT^^2^ positive mice. Injections were performed once per day for 10 days with 2 days break, followed by a washing period for 2 weeks to obtain Drp1^iECKO^ mice.

### Generation of CRISPR/Cas9-engineered mDrp1(C^631^A) knock-in (KI) mutant (mDrp1-KI) mice

The mDrp1-KI mice were generated using Cas9 mRNA (TriLink Bio, L6125100), sgRNA and single-strand DNA homology-directed repair (HDR) method. The gRNA sequence (5’-CCGAGAACAGCGAGATTGTG AGG -3’) was designed using the MIT CRISPR design tool to target the conserved Cys^631^ motif at the C-terminal of mouse mDrp1 gene (corresponding to the Cys^644^ in hDrp1). The single-stranded oligonucleotide incorporating the mDrp1(C^631^A) mutation was used as a template for CRISPR-Cas9 HDR-mediated gene editing in WT mice (C57BL/6). The sgRNA, Cas9 mRNA and single-stranded oligonucleotide encoding mDrp1C^631^A were then injected into the cytoplasm of mouse zygotes and viable two-cell stage embryos were transferred to pseudopregnant female mice by the transgenic and genomic core facility at Augusta University. We analyzed the founders using a combination of PCR and next-generation amplicon sequencing methods to detect the precise mutations and confirmed successful generation of mDrp1(C^631^A)-KI mice. For genotyping offspring, we designed a PCR strategy that was sensitive enough to differentiate between WT and mDrp1(C^631^A) alleles that differ by 3 nucleotides.

### *In vivo* angiogenesis models

All experiments were conducted at 8-12 weeks old male or female littermates of Drp1^fl/fl^ or Cdh5-Cre^+^ (control, wild type (WT)) and Drp1^ECKO^ or Drp1^iECKO^ mice; or C57Bl6 (WT) and mDrp1-KI mice.

### Mouse retinal angiogenesis model

Eyes from postnatal day 5 (p5) mice were enucleated and fixed in 4% paraformaldehyde for 30 min. Retinas were dissected and permeabilized overnight in PBS containing 1% BSA and 0.5% Triton X-100. The permeabilized retinas were incubated with biotin-conjugated isolectinB4 (IB4) (20μg/ml, Sigma-Aldrich), followed by Alexa Fluor 488– conjugated streptavidin (Invitrogen). After washes, samples were flat-mounted using Vectashield (Vector Labs) mounting medium and imaged under Keyence fluorescence microscope. The total length, number of branch points and tip cells of IB4-positive vessels in the retina were quantified on composite high-magnification images using Image J software, as we reported ^27, 41^.

### Hindlimb ischemia model

Mice were subjected to unilateral hindlimb surgery under anesthesia with intraperitoneal administration of ketamine (87 mg/kg) and xylazine (13 mg/kg) as we reported^4, 6, 42^. Briefly, the left femoral artery was exposed, ligated both proximally and distally using 6-0 silk sutures and the vessels between the ligatures were excised. We measured ischemic (left)/nonischemic (right) limb blood flow ratio using a laser Doppler blood flow (LDBF) analyzer (PeriScan PIM 3 System; Perimed).

### Bone marrow transplantation (BMT)

BMT was performed as we previously reported^38, 42^. BM cells were isolated by density gradient separation. Recipient mice were lethally irradiated with 9.5 Gy and received an intravenous injection of 3 million donor BM cells 24 h after irradiation. As reported before transplantation efficiency has been validated^42^. Hindlimb ischemia was induced at 6 to 8 weeks after BMT.

### Isolation of primary mouse aortic ECs

To isolate ECs from mice, the thoracic aorta was cut into 1 mm rings. Each aortic ring was opened and seeded onto a growth factor-reduced matrix with the endothelium facing down. The segments were cultured in EC growth medium for about 4 days. The endothelial sprouting starts as early as day 2. The segments were then removed, and the cells were cultured continually until they reached confluence. The ECs were harvested using neutral proteinase and cultured in EC growth medium for another two passages before experiments, as we reported ^26, 27^.

### Cell Culture and Transfection

The primary HUVECs (human Umbilical Vein Endothelial Cells) from Lonza (CC-2519, USA) were cultured in EndoGRO (EMD Millipore) supplemented with 5% fetal bovine serum (FBS, Atlanta Biological), and used for experiments until passage 6. Cos1 cells (ATCC) were cultured in DMEM (Gibco-BRL) with 10% FBS. For siRNA transfection, overnight serum-starved (0.5% FBS containing culture media) HUVECs were transfected with siRNAs (20 nM) for siControl (Ambion) or siDrp1 (Sense: ACUAUUGAAGGAACUGCAAAAUAUA[dT][dT]; Antisense: UAUAUUUUGCAGUUCCUUCAAUAGU[dT][dT]) from Sigma using Oligofectamine (Invitrogen, 12252011). For plasmid DNA transfection, cells were transfected with DNA (∼6ug) using polyethylenimine (PEI, Polysciences, USA) as we reported ^27^. Adenovirus was transduced in HUVECs as we reported ^29^.

### Mitochondria structure analysis

Overnight serum-starved (0.5%FBS) HUVEC were transfected with Mito-DsRed vector with the mitochondrial targeting sequence or incubated with MitoTracker Red FM (Invitrogen), followed by VEGF (20ng/ml) stimulation for indicated time. Mitochondrial structure was imaged by confocal microscopy ^38^.

### Immunofluorescence analysis in cultured cells ^28, 38^

Cells were seeded onto coverslips and cultured for at least 2 hr prior to fixing and staining. Thereafter cells were fixed with either 4% paraformaldehyde (cat#15710, Electron Microscopy Sciences) or with 3% PFA + 0.75% Gluteraldehyde (cat#16019, Electron Microscopy Sciences) for 10 min at RT and then permeabilized for 3 min with 0.2% Triton-X 100 (Millipore Sigma) Over-extracted cells were treated with 0.4% Triton-X 100. Cells were then washed three times with DPBS and stained overnight at 4°C with primary antibodies diluted in PBS. Next, they were washed twice with PBS for 5 min, incubated with secondary antibodies (diluted 1:1000) for 2 hr at RT in PBS. The following antibodies were used: Rabbit anti-Tom 20 (cat# 11802-1-AP, Proteintech). Preabsorbed secondary antibodies used were anti-mouse IgG 647, anti-rabbit IgG 568 at a 1:1000 dilution. Coverslips were mounted onto slides using ProLong Diamond (Thermo Fisher). All slides were cured at RT in the dark for 2 days prior to imaging.

### Quantification of mitochondria morphology

Cells were plated onto laminin-coated coverslips for 90 min, fixed and stained overnight for Tom20 to label mitochondria. Cells were imaged using the Nikon CSU-W1 spinning disk confocal microscope with a 100X 1.49NA SR objective. Identical image acquisition settings were applied to all experimental groups ^28^. Confocal z-stacks were denoised using Denoise.ai and deconvolved using the Blind algorithm at 20 iterations using NIS-Elements software. This combination of algorithm and number of iterations was chosen because it yielded the highest signal to noise without being destructive to the morphology of Tom20-stained mitochondria. 3D binary objects were created from the deconvolved z-stacks using intensity-based thresholding. The minimum threshold intensity value was manually chosen for each image, and a clean filter was applied to remove punctate background immunofluorescence from the 3D binary mask.

### Modified Boyden Chamber Migration Assay

The upper insert (8-μm pores coated with 0.1% gelatin) containing serum starved HUVEC suspensions (6×10^4^ cells) were placed in the bottom 24-well chamber containing fresh media with 0.5 % FBS and stimulants. The chamber was incubated at 37°C for 6 hr. The upper insert membrane was fixed with 4% PFA for 10 min and stained with crystal violet. Cells remaining on the top of the transwells were removed with cotton swabs and migrated cells were imaged at six random fields and counted as we reported ^27, 41^.

### Cell proliferation assay

The siRNA transfected cells (100,000 cells) were plated on coverslips in 0.5% FBS EndoGro for 24 h to synchronize cells, followed by incubation with 5 µM Bromodeoxyuridine (BrdU) with or without VEGF (20 ng/mL) for 24 h. The BrdU incorporation was detected in fixed, permeabilized and 2N HCl-treated cells followed by staining with anti-BrdU antibody. The cells were taken photos with fluorescence microscope (Keyence, BZ-X700) using ×40 objective. The percentage of BrdU-labelled cells was determined by counting as BrdU-positive/total nuclei as we reported ^27^.

### Fibrin bead *angiogenesis* assay

The capillary-like sprouting fibrin bead assay was performed with minor modifications^40^. HUVEC and fibroblasts (ATCC) were exposed to M199 medium with 10% FBS. HUVEC were attached to dextran-coated Cytodex 3 beads (GE Healthcare Bio-Sciences) and embedded at 250 beads per well in a 24-well plate in a fibrin clot. Fibroblasts were then seeded as a monolayer over the clot at 1.5×10^5^ cells per well. Clots were cultured under standard conditions in EGM-2 medium and medium was changed every day for 5-7 d. Bright-field and fluorescent images were taken, and the sprout number was quantified. The number of sprouts was measured as the number of independent sprouts extruding from each bead and normalized to the number of beads counted, as we reported ^27^.

### Wound scratch cell migration Assay

Confluent ECs were scraped using sterilized 10-μL pipette tips, washed with 0.1% serum media, and stimulated with 20 ng/ml of VEGF for 16 h, Images were captured immediately at 0 hr and at 16 hr after the wounding, as we reported ^43^.

### DCP-Bio1 assay to detect Cys-OH formed (sulfenylated) proteins

To measure sulfenic acid (CysOH) formation (sulfenylation) of proteins, cells were lysed in degassed-specific lysis buffer [50 mM HEPES, pH7.0 at room temperature, 5 mM EDTA, 50 mM NaCl, 50 mM NaF, 1 mM Na_3_VO_4_, 10 mM sodium pyrophosphate, 5 mM Iodoacetamide (IAA), 100 µM DTPA, 1% Triton-X-100, protease inhibitor, 200 unit/mL catalase (Calbiochem), 200 µM DCP-Bio1 (KaraFast, USA)] and then DCP-Bio1-bound proteins were pulled down with streptavidin beads (Thermo scientific, USA) overnight at 4 °C. DCP-Bio1-bound CysOH formed, sulfenylated-proteins were determined by immunoblotting with specific antibodies, as reported ^8, 38, 44^.

### <u>Bi</u>molecular <u>F</u>luorescence <u>C</u>omplementation (BiFC) Assay

COS1 cells were transfected with DNAs (each 250∼500 ng) including N-terminal venus (VN)-flag-AMPK and C-terminal venus (VC)-peptide (negative control) or VC-HA-Drp1-WT or VC-HA-Drp1-C^644^A using PEI for 24 hr. The positive YFP signals by interaction between Drp1 and AMPK were taken by confocal microscopy, as we reported ^27, 38^. pECFP was used as a positive control for transfection.

### Intracellular Redox (ROS) Measurement

To detect intracellular redox state (ROS), HUVECs were transfected with control siRNA and Drp1 siRNA for 48 h. After serum starvation for overnight, cells were stimulated with VEGF (20 ng/ml) for 30 min and then incubated with 5 μM CellRox (Invitrogen) for 30 min at 37°C. Fluorescence signal was measured by confocal microscopy (Zeiss LSM710) using the same exposure condition in each experiment. Relative fluorescence intensity was analyzed using LSM software, as we reported^42^. We confirmed that VEGF-induced increase in CellROX fluorescence was abolished by PEG-catalase.

### MitoSOX (mitochondrial ROS) Measurement

HUVECs were plated on 35 mm glass bottom dishes and then serum-starved (0.5% FBS) for 3 hr, followed by stimulated with VEGF (20 ng/ml) for 15min. Cells were incubated with 5 μM MitoSOX (M36008) for 10 min at 37°C. Fluorescence signals were measured by confocal microscopy with x63 objective and relative fluorescence intensity was calculated using Image J. We confirmed that VEGF-induced increase in MitoSOX fluorescence was abolished by Adeno-Mito-catalase, as we reported ^29^.

### Seahorse XFe24 glycolysis stress test

HUVECs were seeded in the Seahorse XF24 cell culture plate (Agilent, Santa Clara, CA). Wells coated with 0.2% gelatin were used for background subtraction. Cells were washed with Seahorse media supplemented with Glutamine then Seahorse XF24 glycolysis stress test (Agilent, Catalogue # 103020-100, Santa Clara, CA) was employed to evaluate glycolytic capacity according to manufacturer’s instructions, as we reported ^45^.

### Immunofluorescence

Immunofluorescence (IF) staining was performed in fresh frozen 5 μ m thick OCT tissue sections that were stained with primary anti-mouse CD31 antibody (BiosciencesBD Pharmingen™ Purified Rat Anti-Mouse CD31) (1:300). Secondary antibodies were Alexa Fluor 488 or 546-conjugated goat anti-rabbit IgG and goat anti-mouse IgG (Invitrogen)(1:500). Tissue sections were mounted using Vectashield (Vector Laboratories) containing DAPI for nuclear counter-staining. Images were taken using a fluorescence microscope (Keyence, BZ-X700).

### Western blot analysis and Immunoprecipitation

Cells were lysed in buffer [50 mM HEPES (pH 7.4), 5 mM EDTA, 100 mM NaCl, 1% Triton X-100, protease inhibitors (10 μg/ml aprotinin, 1 mmol/L phenylmethylsulfonyl fluoride, 10 μg/ml leupeptin) and phosphatase inhibitors (50 mmol/L sodium fluoride, 1 mmol/L sodium orthovanadate, 10 mmol/L sodium pyrophosphate)]. Lysates, both with or without immunoprecipitation (IP), were used for Western immunoblotting (IB), as we reported^43, 44^.

### Antibodies

Anti-Drp1 (C-5): sc-271583, (IF: 1:200, IB: 1:500, CD31 (BiosciencesBD Pharmingen™ Purified Rat Anti-Mouse CD31) (IF: 1:300), VEGFR2 (CS, 2479, 1:1,000), anti-p-VEGFR2(1175) (CS, 2478, 1:1,000), p-ERK1/2 (CS, 9101, 1:1,000), ERK1/2 (CS, 9102, 1:1,000), p-p38 (CS, 9211, 1:1,000), p38MAPK (Santa Cruz, 7972, 1:1,000), actin (Santa Cruz, 47778, 1:1,000), AMPKα (CS,2532, IP: 1:200, 1B: 1:1000), p-AMPKα (Thr172)(40H9) (CS, 2535, 1:1000), Flag (Sigma-Aldrich, F7425, 3165, 1:1,000), HA (Origene, CB051, TA180128, 1:1,000), Phospho-Acetyl-CoA Carboxylase (p-ACC) (Ser79) (CS,3661, 1:1000), ACC1 (Protein tech 67373-1-Ig, 1:1000), GAPDH (GT239, 1:1000, VDAC (D73D12) CS, 4661, 1:1000).

### Mitochondria and Cytosolic Fractionation

Mitochondria and Cytosol of HUVECs was fractionated by using Mitochondria/Cytosol Fractionation Kit (ab65320).

### Statistical Analysis

Data are presented as mean ± SEM. We performed blinded to group allocation during data collection and analysis. Data were compared between groups of cells and animals by unpaired two tailed Student *t*-test when one comparison was performed or by ANOVA for multiple comparisons. When significance was indicated by ANOVA, the Tukey post-hoc and Bonferroni multiple comparison analysis was used to specify between group differences. Values of *p<0.05, **p<0.01, ***p<0.001were considered statically significant. Statistical tests were performed using Graph pad Prism v8 (GraphPad Software, San Diego, CA).

## Acknowledgements

This work was supported by NIHR01HL160014 (to M.U.-F.), R01HL174014 (to T.F., M.U-F., V.S.), R01HL147550, R01HL133613 (to T.F., M.U.-F.), P01HL160557 (to T.F, M.U-F, E.B.C, D.JR.F, R.L, D.S.), R01HL147639, R01HL155265 (to EJ.B.d.C). NIH VDI P30EY031631 (to R.B.C., T.F., M.U.-F.). VA Merit Review grant I01BX001232 (to T.F.), AHA22TPA971863 (to T.F.), 23TPA107256 (to R.L.), 19EIA34760167 (to EJ.B.d.C), 23POST1022971 (to S.N.), 22CDA (to D.A.), 24CDA (to A.D.), We acknowledge Dr. Sesaki Hiromi at John’s Hopkins University for providing Drp1 floxed mice and Dr. Ralf Adams at Max Planck Institute, Germany for providing Cdh5-Cre^ERT2^ mice. In addition, we acknowledge Dr. Lin Gan at Transgenic & Genomic Editing Core at Augusta University for generating Drp1-C^644^A KI mutant mice. We also thank Dr. Jody L. Martin, a Core director at UC Davis Medical Center for generating adenoviruses.

## Author contributions

M.U.-F., T.F., S.N., Y.MK., designed the study; S.N., Y.MK., A.D., D.A., E.V., V.S., M.S.H., S.Y., S.K., performed research; M.M., S.K., performed genotyping of mice; M.U.-F., T.F., S.N., Y.MK., A.D., D.A., E.V., analyzed data; R.L., D.S., EJ.B.d.C., R.B.C., D.J.F., discussed data and provided inputs; M.U.-F., T.F., Y.NK., S.N., wrote the manuscript; D.J.F., R.L., edited the manuscript. All the authors reviewed the manuscript.

## Supplementary Figures

**Figure S1.**
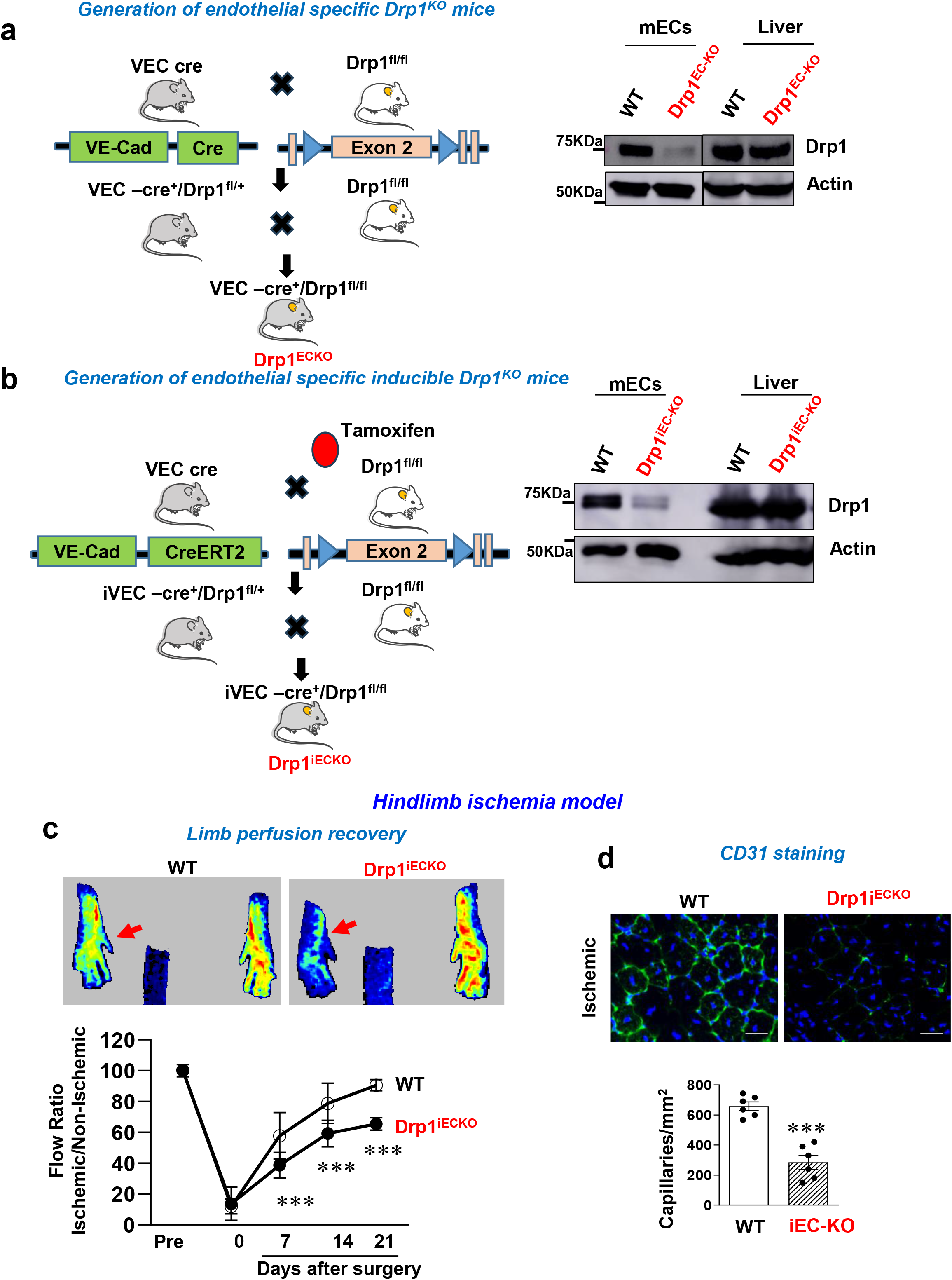
Generation of conditional or inducible EC specific Drp1^KO^ mice. **a and b.** Generation of conditional EC-specific Drp1 KO (Drp1^ECKO^) mice by crossing Drp1^fl/fl^ mice with VE-Cadherin (Cdh5)-Cre mice (**a**) or tamoxifen-inducible EC-specific Drp1 KO (Drp1^iECKO^ Drp1^iECKO^) mice by crossing Drp1^fl/fl^ mice with Cdh5-ERT2-Cre mice which specifically express Cre in ECs upon tamoxifen administration (**b**). Right panels: Lysates from aortic ECs or livers isolated from WT or Drp1^ECKO^ (**a**) or Drp1^iECKO^ (**b**) mice were used for western blotting using Drp1 antibody (Ab) or Actin Ab (loading control). We used Drp1^fl/fl^ mice or Cdh5-Cre^+^ mice as control (WT) in this study and confirmed that both mice showed identical results. **c.** WT or Drp1^iECKO^ mice were subjected to HLI. Left: Limb perfusion recovery determined by the ratio of foot perfusion between ischemic (left) and non-ischemic (right) legs using a laser Doppler flow analyzer. Upper panels show representative laser doppler images of leges on day 21. Right: CD31^+^ capillary density in ischemic gastrocnemius (GC) muscle was measured on day 14 after HLI. Data are mean ± SEM (n=6). ***p<0.001.

**Figure S2.**
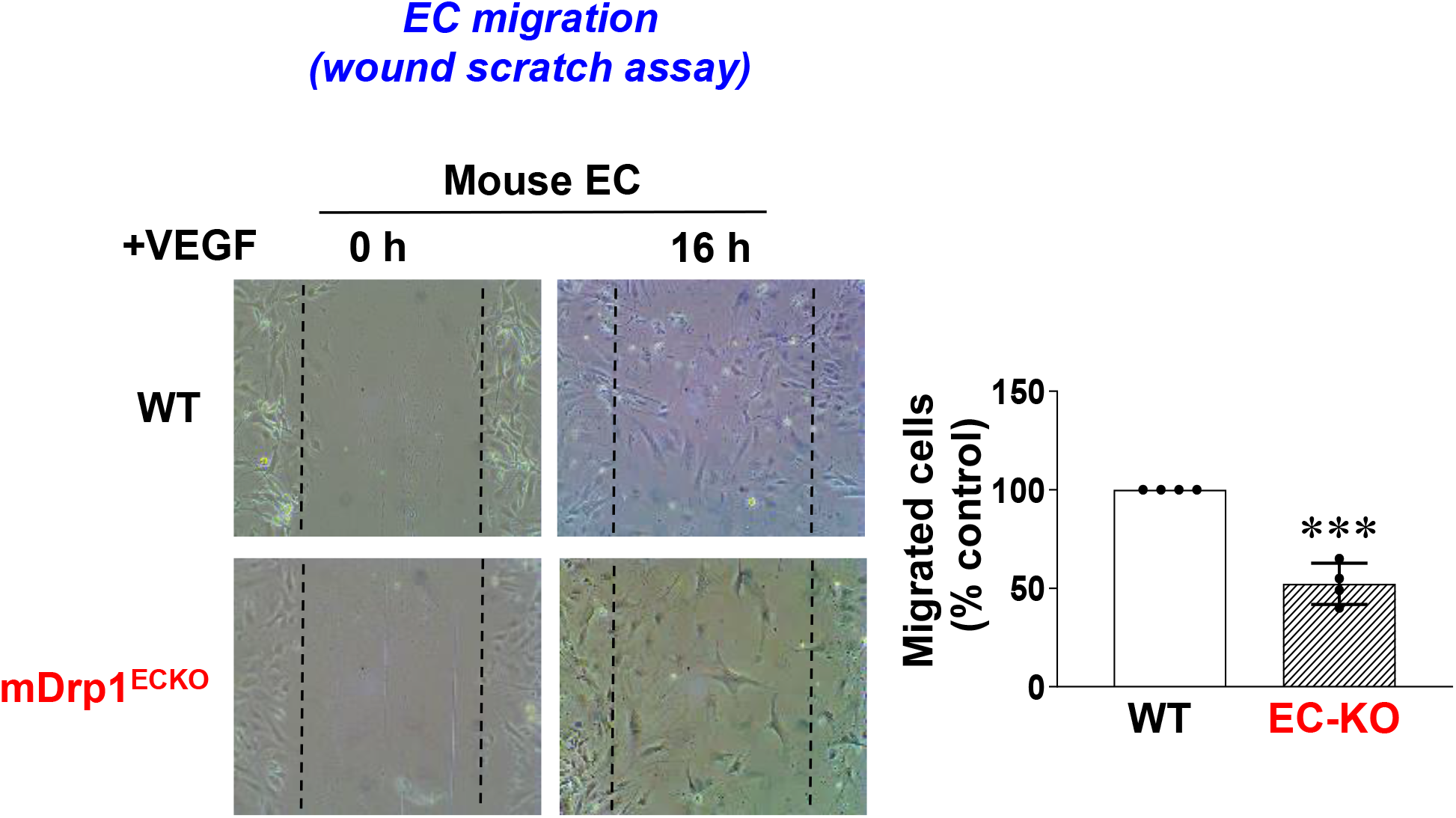
Drp1 is required for would scratch-induced EC migration. **a.** Aortic ECs isolated from WT and Drp1^ECKO^ mice were used to measure wound scratch-induced migration in the presence and absence of VEGF (20ng/ml). Left: Representative images of confluent monolayers of ECs at 0hr and 16 hr after wound scratch. Right graphs represent the averaged % of migrated cells at wounded area in both cells. Data are mean ± SEM (n=4). ***p<0.001.

**Figure S3:**
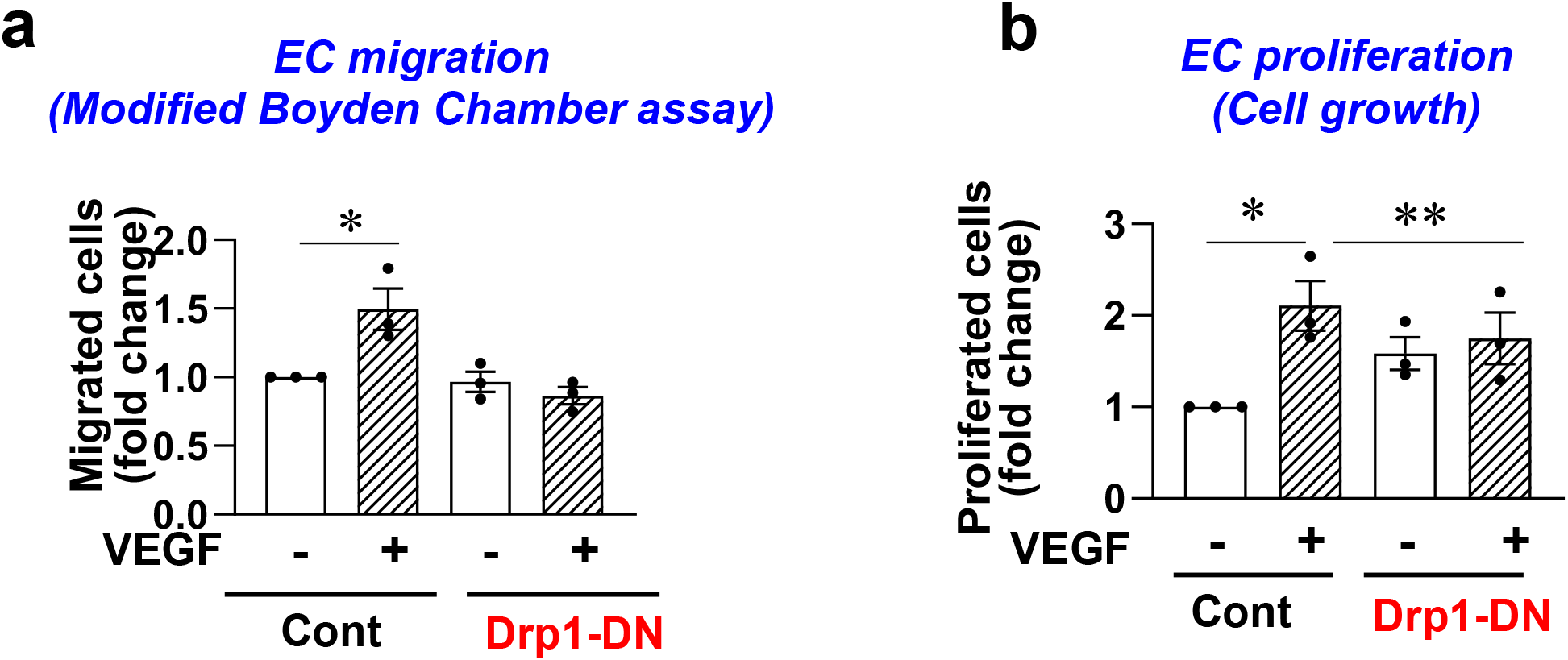
Drp1 GTPase activity is required for VEGF-induced angiogenic responses in ECs. **a and b.** HUVECs transduced with Ad.null (Control) or Ad.Drp1-dominnat negative (DN) mutant were used to measure VEGF (20ng/ml)-induced EC migration (Modified Boyden Chamber assay)**(a)** or EC proliferation (BrdU incorporation)**(b)**. Data are mean ± SEM (n=3). *p<0.05, **p<0.01.

**Figure S4.**
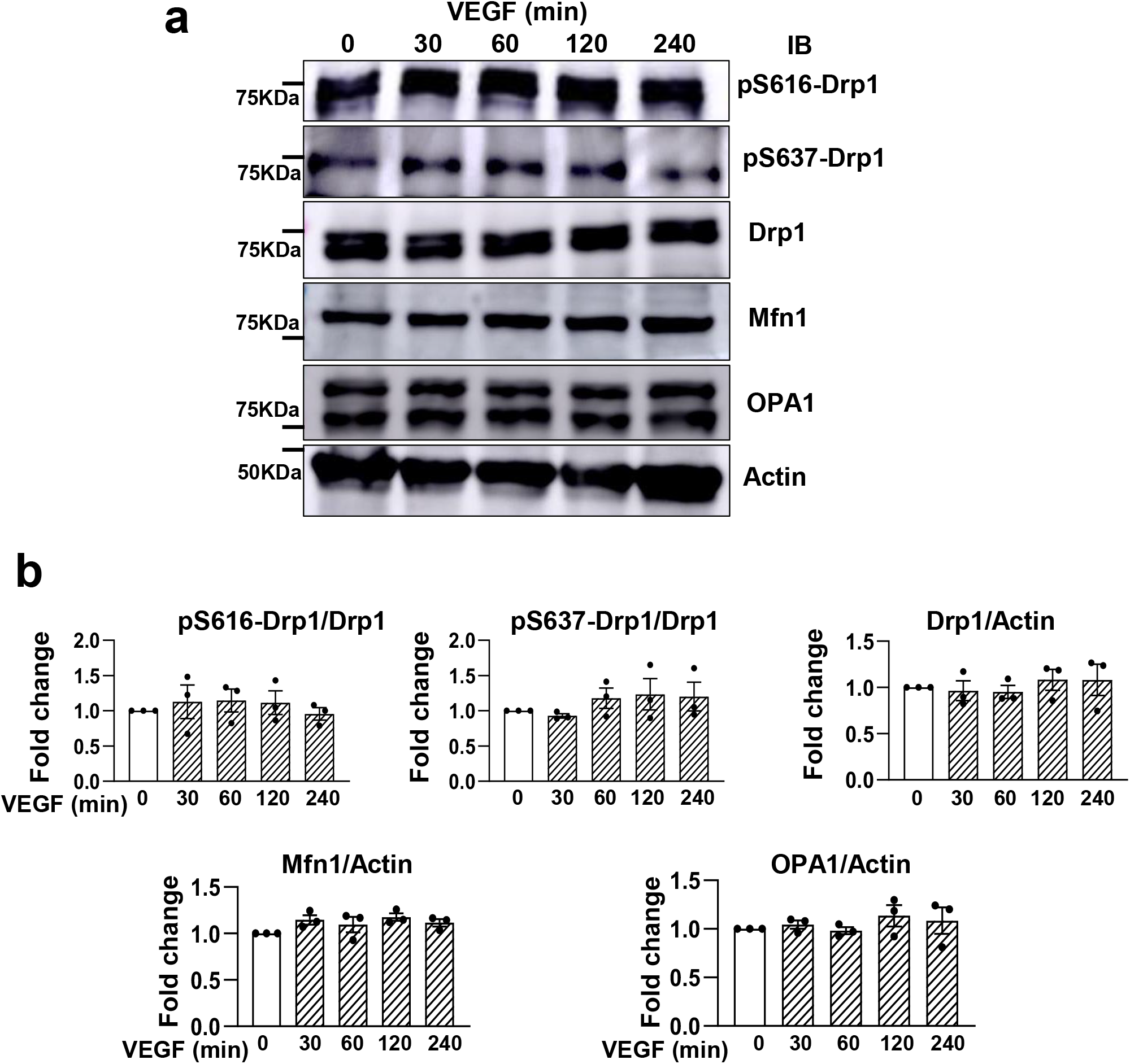
VEGF does not change phosphorylation of Drp1 or total protein expression of mitochondrial dynamics proteins in ECs. **a and b.** HUVECs were stimulated with VEGF (20ng/ml) for the indicated time and total lysates were IB with antibodies indicated (**a**). Bottom panels show averaged fold change from the basal control (**b**).

**Figure S5.**
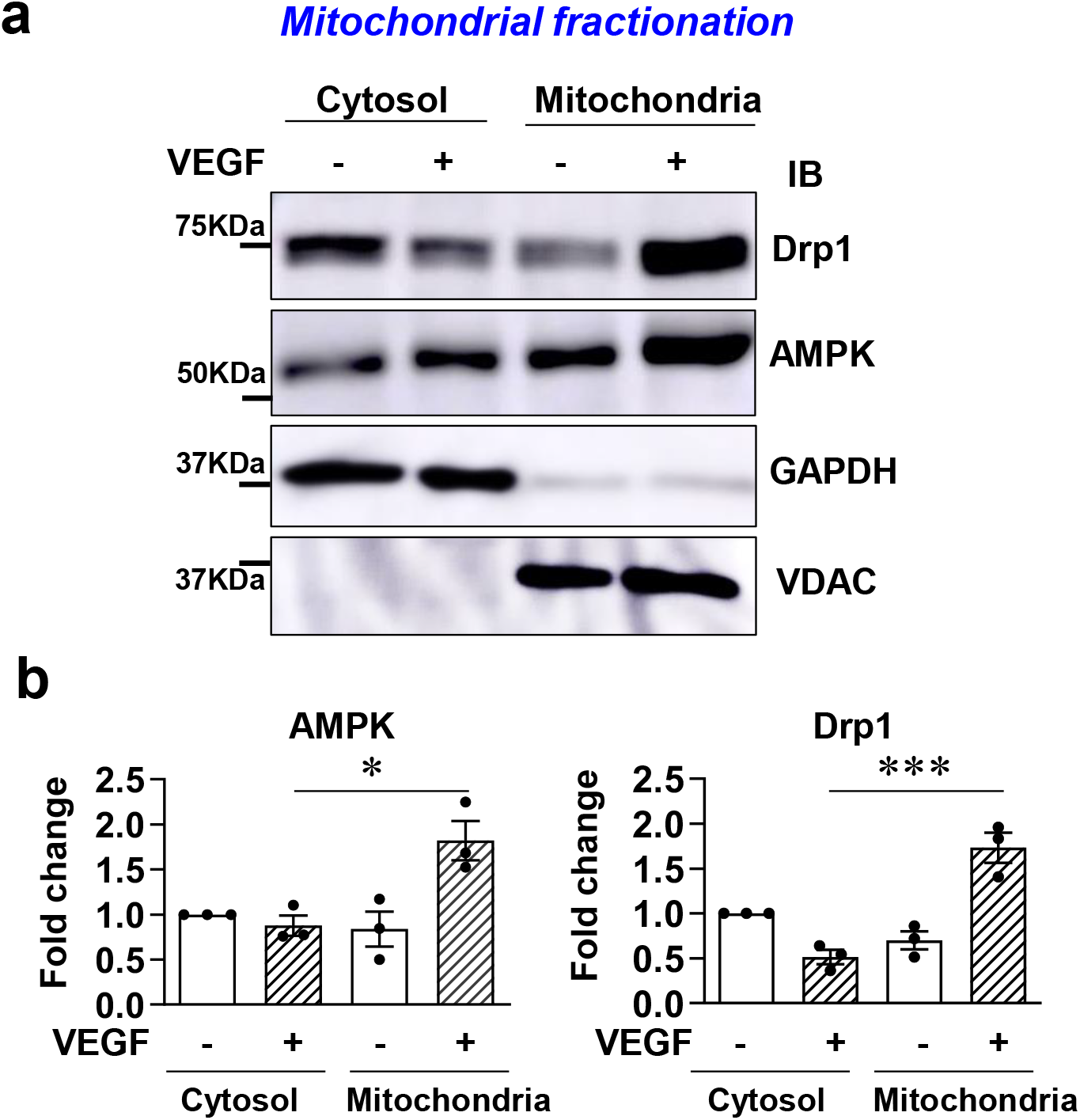
VEGF stimulation increases Drp1 and AMPK protein expression in mitochondrial fraction in ECs. HUVECs stimulated with VEGF for 15 min were used for isolating Mitochondrial and cytosolic fractions. Each fraction was used for IB with anti-Drp1 or - AMPK Ab or -GAPDH (cytosol marker) or -VDAC (mitochondrial marker) Ab. Bottom panels represent fold change from basal in cytosol fraction. Data are mean ± SEM (n=3). *p<0.05, ***p<0.001.

**Figure S6.**
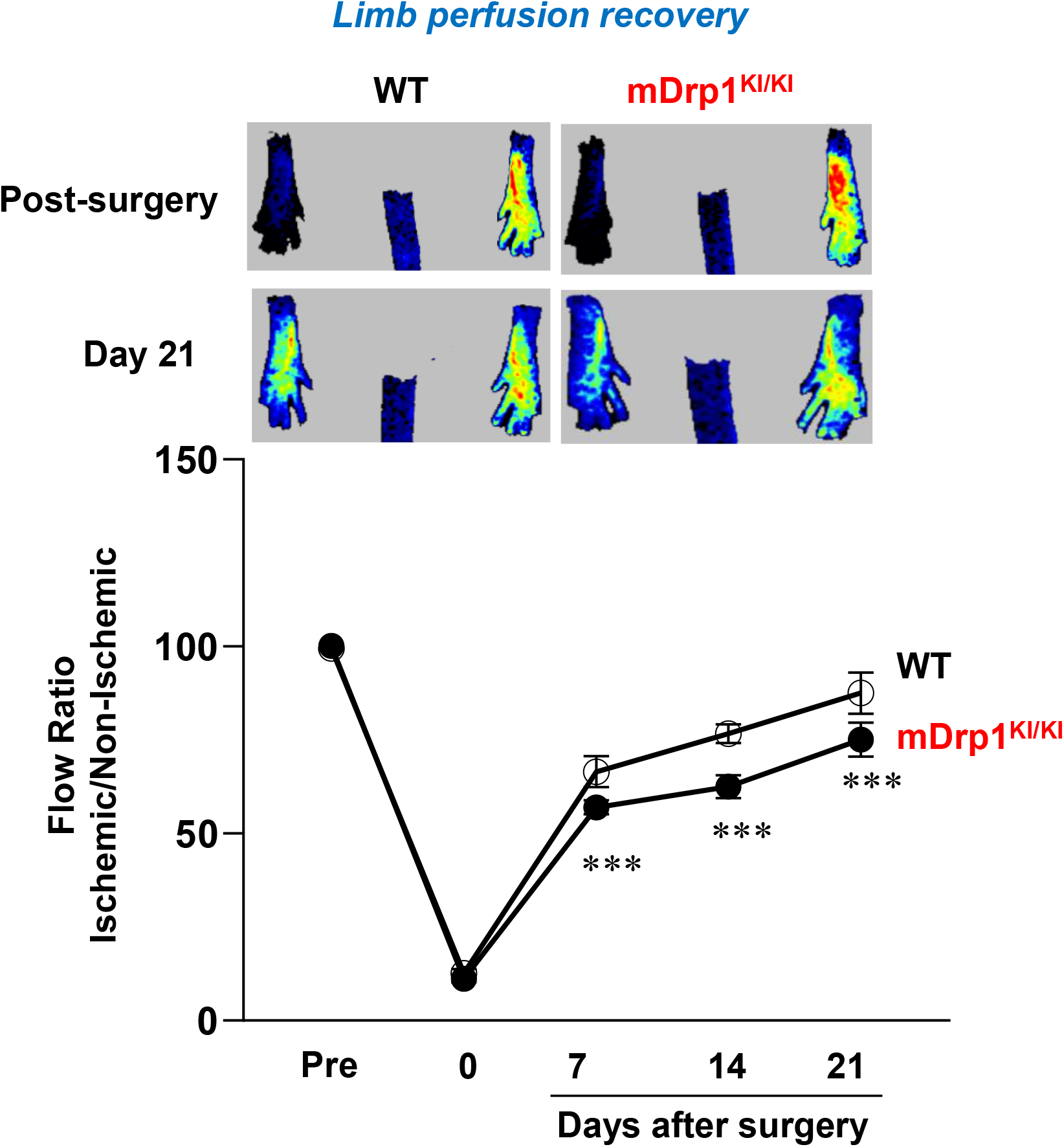
WT or mDrp1^KI/KI^ mice were subjected to HLI and limb perfusion recovery was determined by the ratio of foot perfusion between ischemic and non-ischemic legs after HLI using a laser Doppler flow analyzer. Upper panels show representative laser Doppler images of legs in WT and mDrp1^KI/KI^ mice at post-surgery (day 0) and day 21 after HLI. Data are mean ± SEM (n=6) ***p<0.001.

